# Single molecule analyses of *Salmonella* translocated effector proteins reveal targeting to and dynamics in host cell endomembranes

**DOI:** 10.1101/2022.05.16.492065

**Authors:** Vera Göser, Marc Schulte, Felix Scharte, Rico Franzkoch, Viktoria Liss, Olympia E. Psathaki, Michael Hensel

## Abstract

Bacterial pathogens deliver proteins in temporal and spatial coordinated manner to manipulate mammalian host cells. The facultative intracellular pathogen *Salmonella enterica* remodels the host endosomal system for survival and proliferation inside host cells. The pathogen resides in a membrane-bound compartment termed *Salmonella-*containing vacuole (SCV). By *Salmonella*- induced fusions of host endomembranes, the SCV is connected with extensive tubular structures termed *Salmonella-*induced filaments (SIF). The intracellular lifestyle of *Salmonella* critically depends on effector molecules translocated by the SPI2-encoded type III secretion system (SPI2-T3SS) into host cells. A subset of these effectors is associated with, or integral in SCV and SIF membranes. It remained to be determined how SPI2-T3SS effectors reach their subcellular destination, and how these effectors interact with endomembranes remodeled by *Salmonella*. We deployed self-labeling enzyme (SLE) tags as novel approach to label translocated effector proteins in living host cells, and analyzed their dynamics on single molecule level. We found that SPI2-T3SS effector proteins diffuse in membranes of SIF with mobility comparable to membrane-integral host proteins in endomembranes. Dynamics differed between various effector proteins investigated and was dependent on membrane architecture of SIF. In the early infection, we observed host endosomal vesicles associated with *Salmonella* effector proteins. Effector-positive vesicles continuously fused with SCV and SIF membranes, providing a route of effector delivery by SPI2-T3SS translocation, interaction with endosomal vesicles, and ultimately fusion with the continuum of SCV/SIF membranes. This novel mechanism controls membrane deformation and vesicular fusion to generate the specific intracellular niche for bacterial survival and proliferation.

## Introduction

Various intracellular pathogens are confined to membrane-bound compartments. Within these organelles, pathogens are able to adopt specific intracellular lifestyles. Biogenesis of specialized pathogen-containing vacuoles depends on recruitment of subsets of host cell endosomes in order to establish nutritional supply, and to evade the host immune defense. For this purpose, pathogens translocate, by different secretion systems, specific effector proteins that manipulate the host cell endocytic system (Weber & Faris, 2018).

*Salmonella enterica* is a Gram-negative, foodborne bacterial pathogen, causing diseases ranging from severe typhoid fever to self-limiting gastrointestinal infections in various hosts. *S. enterica* serovar Typhimurium (STM) is commonly used to investigated the intracellular lifestyle of *Salmonella*. After invasion or phagocytic uptake, STM initiates a complex intracellular lifestyle enabling survival and proliferation within host cells. STM resides in a membrane-bound compartment termed *Salmonella-*containing vacuole (SCV), which acquires late endosomal markers, but does not mature to a bactericidal compartment (LaRock et al., 2015). Characteristic for infected cells is the formation of tubular structures connected to the SCV. Such *Salmonella-*induced tubules (SIT) comprise various tubular structures composed of recruited host endomembranes of various organellar origin (Schroeder et al., 2011). The best characterized SIT are *Salmonella-*induced filaments (SIF), marked by lysosomal membrane glycoproteins such as LAMP1 (Garcia-del Portillo et al., 1993). SIF are highly dynamic in the initial phase of intracellular lifestyle. If fully developed, SIF comprise double membranes built up during development where initial SIF are single-membrane tubular compartments (leading SIF), which over time, convert into double-membrane (trailing SIF) tubular structures (Krieger et al., 2014). The molecular mechanisms of these pathogen-driven events of vesicle fusion and membrane deformation remain to be determined.

SIF formation and systemic virulence of STM are dependent on functions of genes within *Salmonella* pathogenicity island 2 (SPI2). SPI2 encodes a type III secretion system (T3SS) which enables the translocation of various effector proteins inside the host cell (Figueira & Holden, 2012). Mutant strains deficient in SPI2-T3SS are highly attenuated in systemic virulence in the mouse model of systemic infection, and show reduced intracellular replication in cell-based models (Hensel et al., 1995; Hensel et al., 1998). We recently reported that SIF biogenesis supports intracellular lifestyle by bypassing nutritional restriction in SCV-SIF continuum by recruiting nutrients from the host endosomal system and is therefore crucial for bacterial proliferation and survival (Liss et al., 2017).

Despite the large number of effector proteins translocated by the SPI2-T3SS, only a subset of these manipulates the host endosomal system and induces vesicle fusion for SCV and SIF biogenesis. These are SifA, SseF, SseG, PipB2, SseJ and SopD2 (Figueira & Holden, 2012). The most severe phenotype is mediated by SifA, as mutant strains defective in *sifA* fail to induce SIF and show loss of SCV integrity leading to bacterial release into host cytosol, and attenuation in intracellular proliferation and systemic virulence (Beuzon et al., 2000). SseF, SseG and PipB2 contribute to SIF formation, as mutant strains lacking the effectors show aberrant SIF biogenesis. Infection with *sseF*-deficient STM leads to the formation of only single-membrane SIF, and infection with *pipB2*-deficient strains results in the induction of enlarged bulky SIF (Krieger et al., 2014; Rajashekar et al., 2014).

A subsets of SPI2-T3SS effectors is recruited to *Salmonella*-modified membranes (SMM) after translocation that are prominently associated with membranes of the SCV and SIF network (Kuhle & Hensel, 2002). This subcellular localization appears to be prerequisite for effector and host protein interactions that mediate vesicle fusion and SIF elongation (Fang & Meresse, 2022). However, the molecular mechanisms of effector targeting to SMM are poorly understood. In STM-infected cells, a dynamic extension of SIF network was observed, raising the question how SPI2-T3SS effector integrate into SMM. For example, highly hydrophobic effectors like SseF appear exclusively associated with SIF membranes (Abrahams et al., 2006; Müller et al., 2012). As T3SS translocation delivers effector proteins into host cell cytosol, specific mechanisms of targeting to and integration into host endosomal membranes are required, and we here applied novel imaging approaches to unravel these mechanisms.

We recently established an imaging approach utilizing self-labeling enzyme (SLE) tags fused to STM effector proteins to enable super-resolution microscopy (SRM), and single molecule imaging of effector dynamics in living cells (Göser et al., 2019). Here we applied these approaches to investigate the delivery of SPI2-T3SS effectors to SMM, their localization and dynamics in SMM. This study provides new insights in the delivery mechanism of effector proteins to the SCV-SIF continuum.

## Results

### Continuous interactions of SCV, SIF and host cell endosomal compartments

We analyzed interactions of intracellular STM with the host cell endosomal system. Pulse/chase experiments with fluorochrome-conjugated gold nanoparticles (nanogold) allowed to label the lumen of endosomes (Zhang & Hensel, 2013). A subset of these endosomes was in contact with dynamic SIF and events of fusion between nanogold-labeled endosomes and SIF were detected (**Movie 1**). Due to the transient nature, fusions between host cell endosomes and membranes of SCV or SIF were rarely determined, and **Movie 1** shows a representative event. In contrast to other fluid tracers that become rapidly diluted after fusion between endosomes and SIF, the aggregation of nanogold led to formation of distinct foci that were readily detectable after fusion events.

Using live-cell correlative light and electron microscopy (CLEM), infected cells were imaged during the formation of dynamically extending SIF (**Fig. 1**). Correlation identified LAMP1-positive tubular vesicles in connection with SCV harboring STM. The investigated cell showed double-membrane (dm) SIF distal to the SCV (**Fig. 1E**), or in connection to SCV (**Fig. 1F, G**). In few occasions, the contact of single-membrane (sm) vesicles of spherical appearance with dm SIF was observed (**Fig. 1H, I, J**). Although the temporal resolution of our live-cell imaging (LCI) approach did not allow to distinguish fusion from fission events for the vesicle, our data would be in line with fusion of a host cell endosome with a dm SIF and release of luminal content in the outer lumen of SIF.

**Fig. 1.**
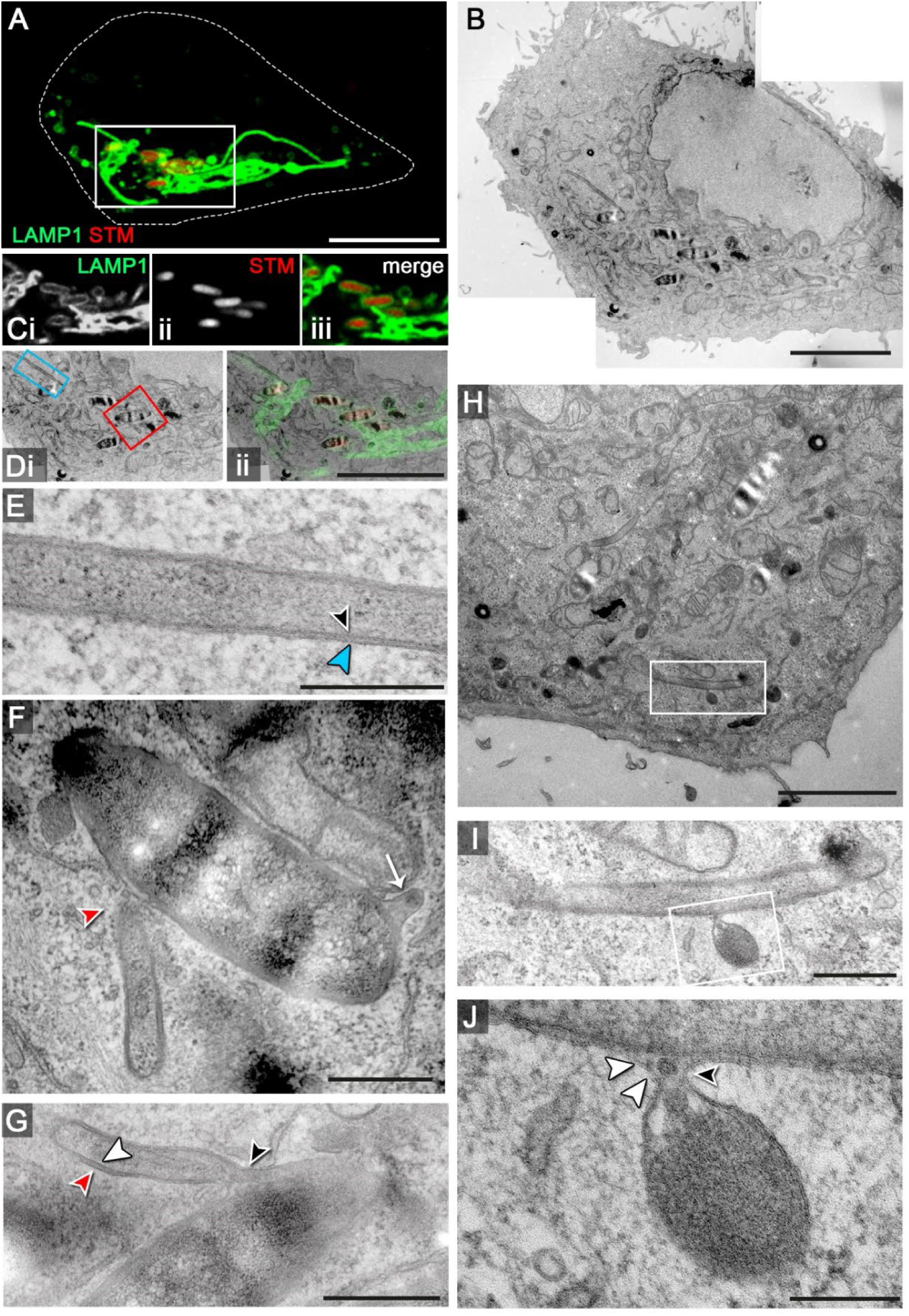
Interactions of host cell endosomal membranes in STM-infected cells. HeLa cells stably expressing LAMP1-GFP (green) were infected with STM WT expressing mCherry (red) and CLSM was performed (**A, C**) to identify SIF-positive cells showing dynamic extension of SIF networks. Cells were fixed 7 h p.i., coordinates registered and samples were processed for TEM (**B**). Correlation of CLSM and TEM modalities allowed identification of STM in SCV and extending SIF tubules (**D**). Regions of interest are indicated by boxes and details show a double-membrane (dm) SIF tubule distal to SCV (blue, **E**), and in connection with the SCV (**F, G**). The white box in panel **H** indicates an event of vesicle interaction with a dm SIF, and details are shown in higher magnification (**I, J**). Arrowhead indicate interactions with double membrane compartments, while single-membrane tubules are indicated by arrows. Scale bars: 10 µm, 5 µm (**B**, **C**), 3 µm (**H**), 500 nm (**E**, **F**, **I**), 300 nm (**G**), 200 nm (**J**).

### Distribution of SPI2-T3SS effector proteins on SCV and SIF membranes

The previous data revealed the fusogenic properties of the SCV/SIF continuum, and indicated how the SIF network is dynamically expanding. Prior work demonstrated that formation of the SIF network is dependent on translocated SPI2-T3SS effector proteins, and that subsets of these effector proteins are closely associated with SIF membranes (Knodler et al., 2003; Miao & Miller, 2000; Müller et al., 2012). Thus, we next followed the distribution of PipB2 as representative SPI2-T3SS effector protein over the course of STM infection (**Fig. 2A**). In the early phase (4 h p.i.), the signal intensity for PipB2 immunostaining was low, and most of the signals were associated with small spherical vesicles. At 8 h p.i., a SIF network was developed and PipB2 signals were mostly associated with SIF and SCV membranes. At the end of observation, i.e. 16 h p.i., PipB2 signal was strongly increased and was almost exclusively colocalized to membranes of SIF and SCV. A similar subcellular distribution over time of infection was observed for other membrane-associated SPI2-T3SS effectors such as SseF and SseJ (**Fig. S 1**).

**Fig. 2.**
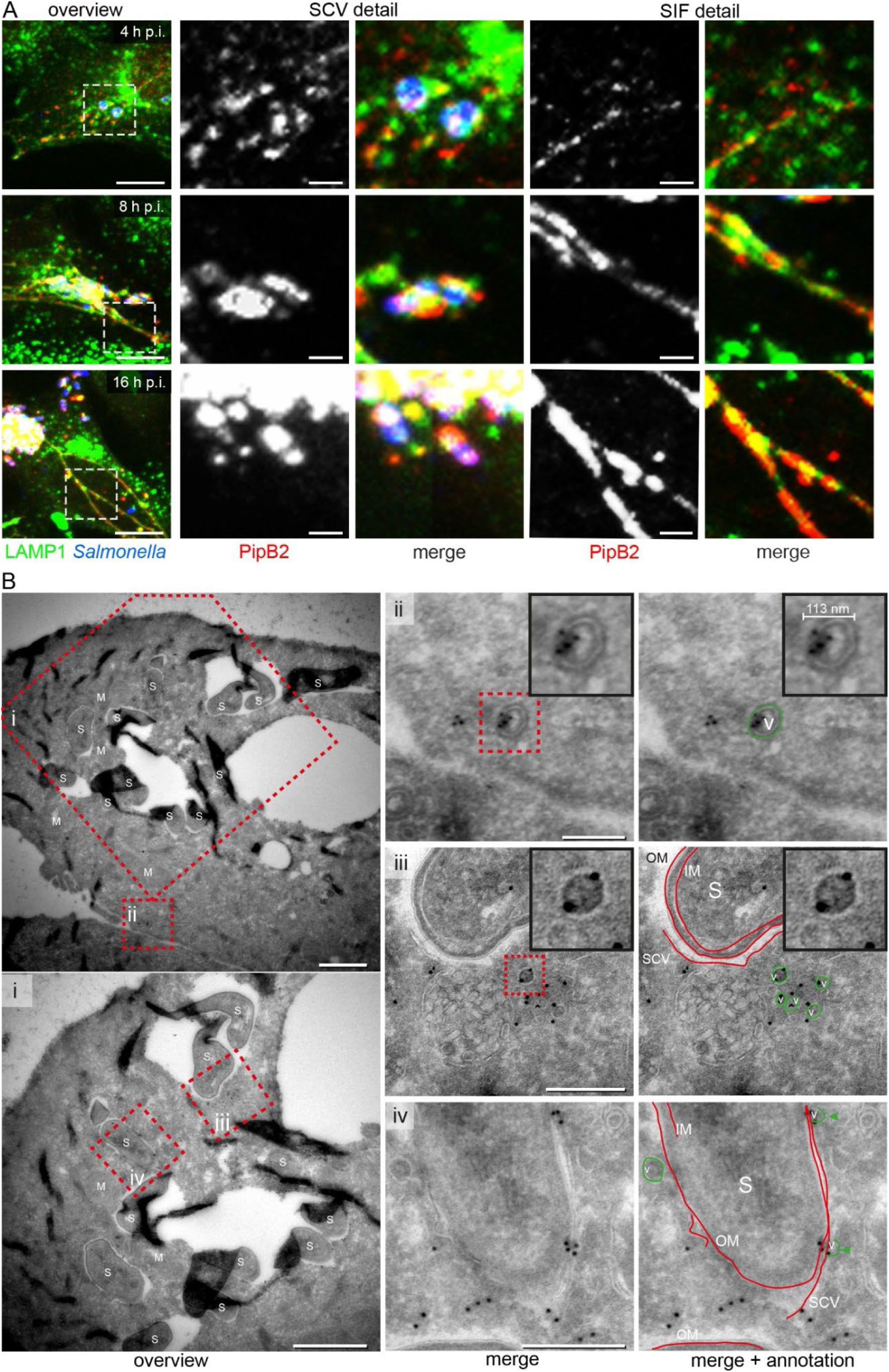
Kinetics of distribution of SPI2-T3SS effector proteins and vesicular localization of translocated SseF. **A)** Distribution of translocated effector proteins over the course of infection. HeLa cells stably expressing LAMP1-GFP (HeLa LAMP1-GFP) were infected with STM WT expressing *pipB2*::M45. At various time points after infection, cells were fixed and immunolabeled for STM (blue) and effector proteins (red). Details of SCV and SIF are shown. Scale bars: 10 and 2 µm in overview and details, respectively. **B)** Vesicular localization of translocated SseF revealed by immunogold EM. HeLa LAMP1-GFP cells were infected with STM WT expressing *sseF*::3xHA and fixed 8 h p.i. The samples were processed for immunogold labeling for HA-tagged SseF. (Details of overviews (**B** and **i**) of SseF immunogold-labeled sections are shown in ii-iv. **ii)** A subset of triple HA-tagged SseF immunogold labeling is associated with the outer and inner side of spherical vesicular membranes. See also color highlighted vesicle structure in green on the left. Inserts strongly clarify localization of immunogold label inside the vesicle and on the vesicular membrane. **iii)** Triple HA-tagged SseF immunogold labeling is also found on endomembranes mostly in close proximity to vesicles. Color marking in green for vesicles and red for SCV, inner (IM) and outer (OM) bacterial membrane is highlighting the distribution of gold labeling on membrane structures. Inserts strongly clarify localization of immunogold label on the vesicular membrane. **iv)** The majority of triple HA-tagged SseF immunogold labeling is distributed on endomembranes, specifically on membranes closely associated with the SCV and directly on the SCV. See also color marking in red indicating for SCV membrane, IM and OM. Scale bars: 1 µm in overviews (B, i); 250 nm in ii, iii, and iv.

To investigate if translocated effector proteins are present in membranes of endosomal compartments prior to integration of these membranes into the SCV/SIF continuum, we applied immunogold labeling for TEM analyses. A triple HA-tagged allele of *sseF* was used as representative membrane-integral SPI2-T3SS effector protein. SseF-3xHA was synthesized, translocated and subcellular localized as observed for SseF-HA (**Fig. S 2**). For highest preservation of endosomal membranes and epitopes for immunolabeling, the Tokuyasu technique (Tokuyasu, 1973) was applied. In infected HeLa cells, immunogold-labeled SseF-3xHA was associated with SCV membranes (**Fig. 2**). We also detected labeling for SseF associated with spherical membranes compartments distal to the SCV. In ultrathin sections, such signal could result from cross-sections of small spherical vesicles, or of extended tubular compartments such as SIF. To distinguish these forms, consecutive ultrathin sections were inspected, indicating labeling a single section rather than in compartments extended through several sections (**Fig. S 2**).

These ultrastructural observations support a model that effector proteins associate with and integrate in host cell endosomal membranes prior to incorporation into the SCV/SIF continuum.

### Models for SPI2-T3SS effector targeting to endomembranes

It is not known how hydrophobic effector proteins insert into host cell endomembranes. We built several hypotheses for the route of SPI2-T3SS effector proteins from translocation to their final destination (**Fig. 3**). In model A, effector proteins are directly integrated into SCV membranes after translocation. In model B, effector proteins are translocated into the host cell cytosol, and a fast interaction with unknown bacterial or host cell proteins enables insertion into host endomembranes. In model C, direct delivery of effector proteins into host vesicular membranes is mediated by the SPI2-T3SS itself, and no cytosolic effector intermediates are present. In model A, peripheral distribution of effector proteins is mediated by tubulation of SCV membranes containing effector proteins. In models B and C, effector proteins are first inserted into endosomal membranes that subsequently fuse with developing SIF. We would also consider combinations of the models, and distinct modes of delivery for different effector proteins. We set out to test these models by applying a recently developed LCI approach for translocated effector proteins on single molecule level (Göser et al., 2019).

**Fig. 3.**
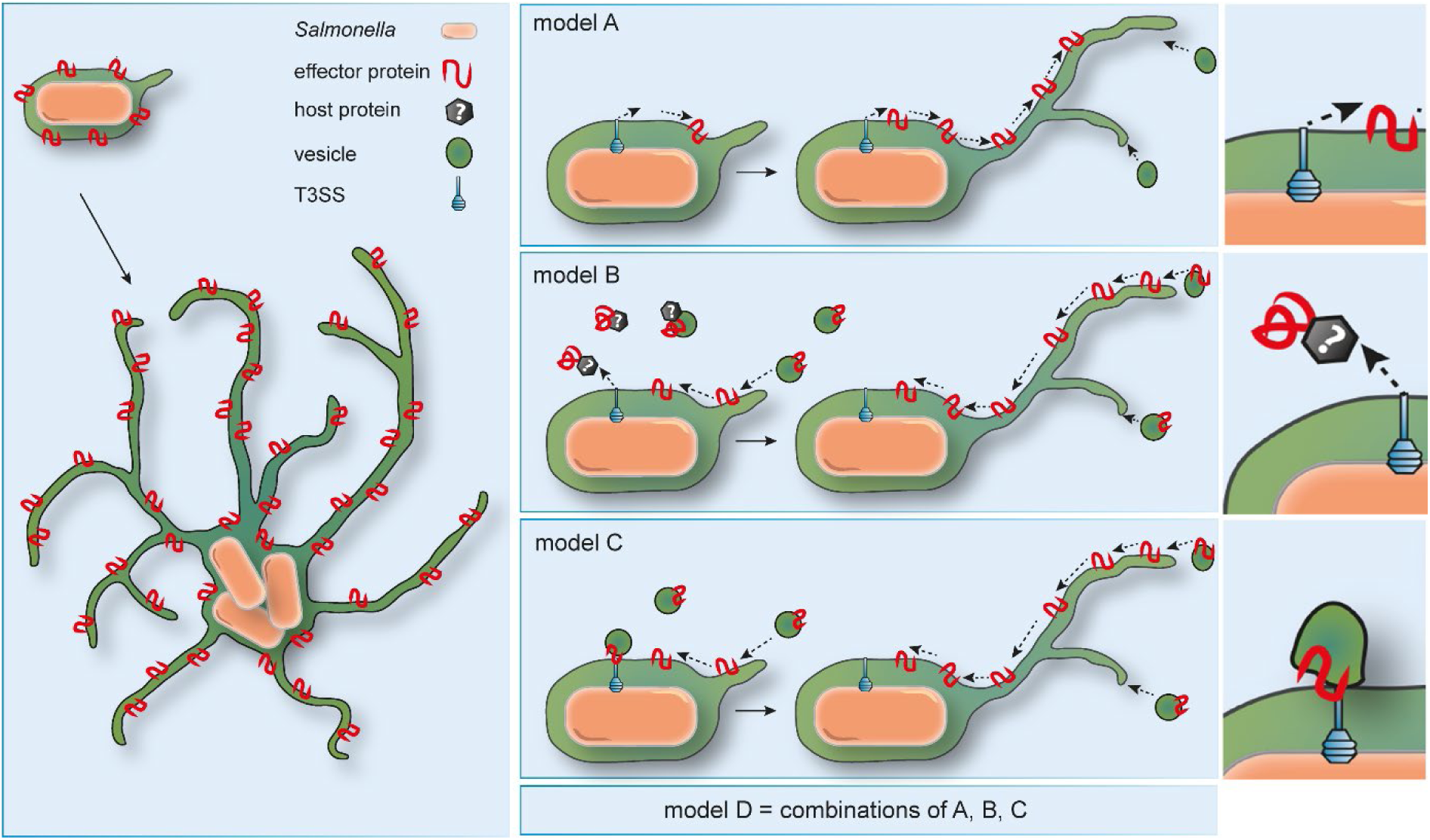
Models for targeting of SPI2-T3SS effector proteins to host cell endosomal membranes. Model A: Effector proteins are directly inserted into the SCV membrane after translocation and diffuse from the SCV to the periphery of SIF. Model B: Effector proteins are translocated into the cytosol, chaperoned and inserted into endomembranes by unknown host factors. Effector proteins are delivered by fusion of host endocytic vesicles with SIF and SCV. Model C: Endomembranes are recruited to SCV and T3SS where effector proteins are directly delivered and inserted into endomembranes. After fusion of endocytic vesicles effector proteins are delivered. Combinations of models A, B, and C may be considered.

### SPI2-T3SS effector proteins are highly dynamic on SIF membranes

To follow the dynamics of SPI2-T3SS effector proteins on or in SIF membranes, we deployed single molecule localization and tracking microscopy (TALM) (Appelhans et al., 2018). As host cells, HeLa cells were used that constitutively express LAMP1-monomeric enhanced green fluorescent protein (LAMP1-GFP) to allow visualization of SCV and SIF. Host cells were infected with STM mutant strains deficient in genes for specific effectors. The strains harbored plasmids encoding effector proteins fused to HaloTag, a self-labeling enzyme (SLE) tag, and infected cells were labeled with HaloTag ligand coupled to the fluorescent dye tetramethylrhodamine (HTL-TMR). As previously shown (Göser et al., 2019), the effector proteins SseF, SifA and PipB2 fused to HaloTag can be localized in infected host cells 8 h p.i. and a complete colocalization with LAMP1-GFP-positive SCV and SIF membranes was observed (**Fig. 4A, Fig. S 3, Movie 2, Movie 4, Movie 5**).

**Fig. 4.**
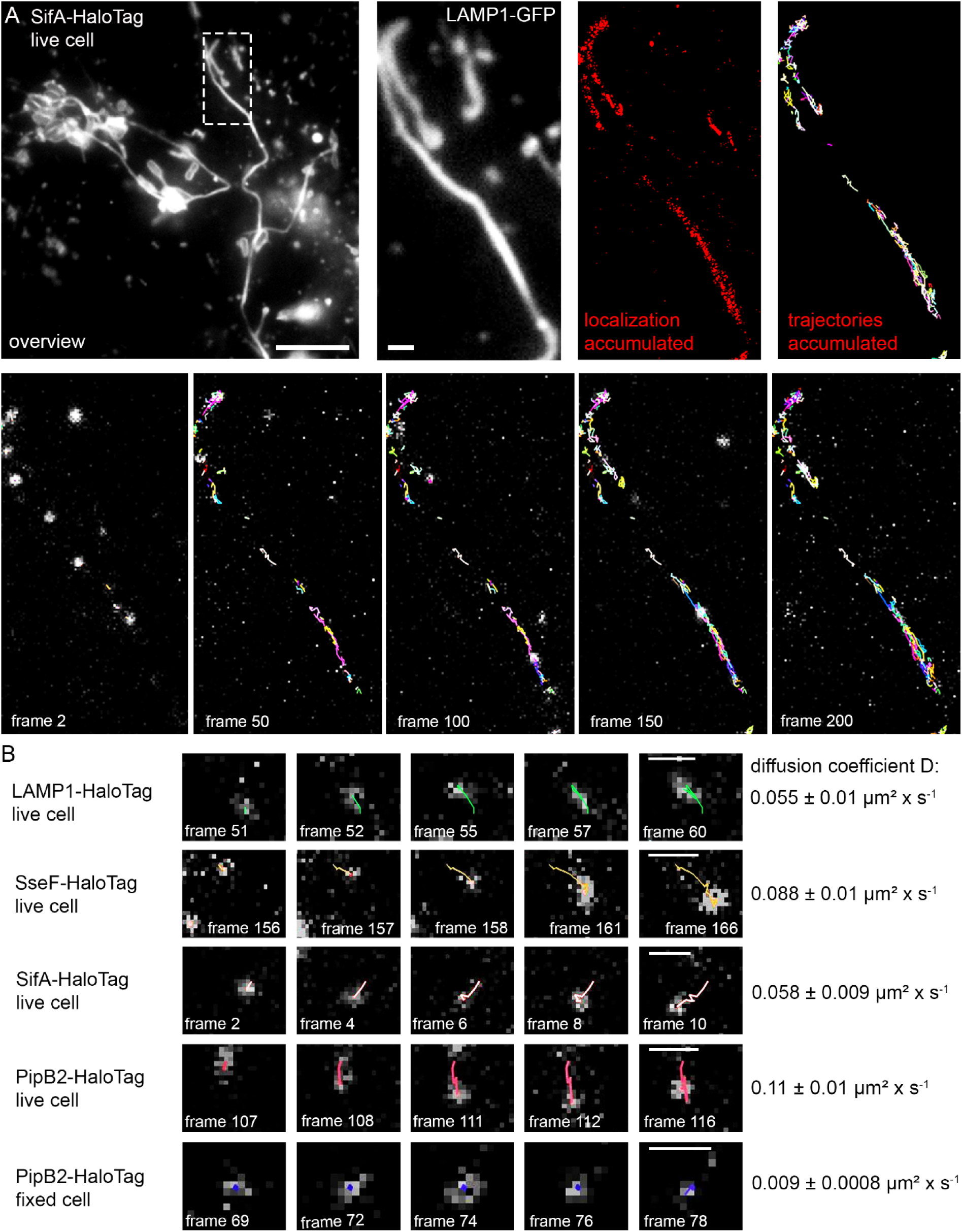
Single molecule localization and tracking of STM SPI2-T3SS effector proteins on double-membrane SIF. HeLa cells stably expressing LAMP1-GFP were infected with STM *sifA* mutant strain expressing SifA-HaloTag with a multiplicity of infection (MOI) of 75. Following incubation for 7 h under standard cell culture conditions, LCI was performed. Labeling reactions were performed directly bevor imaging, using HTL-TMR with a final concentration of 20 nM for 15 min at 37 °C. **A)** Shown are representative SRM images acquired using 15% laser power at the focal plane, rendered from single molecule localizations (SML) and tracking (SMT) within 200 consecutive frames. Selected frames (frame rate: 32 frames per s) of the TMR signal, localization and tracking are presented (also showing elapsed trajectories). **B)** Selected frames of trajectories from a single molecule. Using at least 2,800 pooled trajectories for proteins in at least 20 infected cells in three biological replicates recorded under the same conditions, the diffusion coefficient D was calculated using the Jaqaman algorithm. The indicated error represents the calculated error of the resulting slope (with 95% confidence bounds). Scale bars: 10 and 1 µm in overviews and details, respectively. A sequence of 200 frames for SifA-HaloTag is shown in **Movie 2**.

We tracked the movement of the key STM effector proteins SseF, SifA and PipB2 fused to HaloTag on *Salmonella*-modified membranes. By analyzing comprehensive data sets of single molecule trajectories, the mobility of effector proteins using pooled trajectories resulting in two-dimensional diffusion coefficient (DC), extracted from mean square displacements (MSD), was demonstrated (Göser et al., 2019). As control, the host membrane protein LAMP1 was used and visualized after transient transfection of HeLa LAMP1-GFP cells for expression of LAMP1-HaloTag (**Movie 3**). For non-moving particles, tracking of PipB2-HaloTag on SIF tubules was performed in fixed host cells (**Movie 6**).

The DC of fixed PipB2-HaloTag was quantified as 0.009 +/- 0.0008 µm^2^ x s^-1^. For LAMP1- HaloTag, a DC of 0.11 +/- 0.01 µm^2^ x s^-1^ was determined. The effector proteins SifA, SseF, and PipB2 fused to HaloTag varied in their mobility with DC of 0.058 +/- 0.009, 0.088 +/- 0.01 and 0.11 +/- 0.01 µm^2^ x s^-1^, respectively (**Fig. 4B**, and **Movie 2**, **Movie 4**, **Movie 5**). In all cases, the trajectories developed bidirectional, without preferential movement of molecules towards SCV- proximal or SCV-peripheral portions of SIF.

Infection with STM mutant strain Δ*sseF* leads to increased formation of sm SIF which are smaller in diameter and volume. Fully developed SIF in STM WT-infected cells are predominantly dm SIF (Krieger *et al*., 2014; Rajashekar *et al*., 2014). We analyzed SPI2-T3SS effector mobility on sm SIF to investigate potential effects of SIF architecture on distribution and diffusion of effector proteins. HeLa LAMP1-GFP cells with STM Δ*sifA* Δ*sseF* strain expressing *sifA*::HaloTag and dynamics of SifA-HaloTag molecules on sm SIF were analyzed (**Fig. 5AB**, **Movie 7**, **Movie 8**). The reduced diameters of sm SIF were verified by intensity profile analyses of accumulated SifA-HaloTag trajectories on SIF induced by STM WT and Δ*sseF* strains (**Fig. 5D**). When calculating DCs for LAMP1-HaloTag and SifA-HaloTag on sm SIF induced in cells infected by STN Δ*sseF*, a reduction of mobility with DC values of 0.028 +/- 0.008 and 0.035 +/- 0.004 µm^2^ x s^-1^, respectively was observed (**Fig. 5C**).

**Fig. 5.**
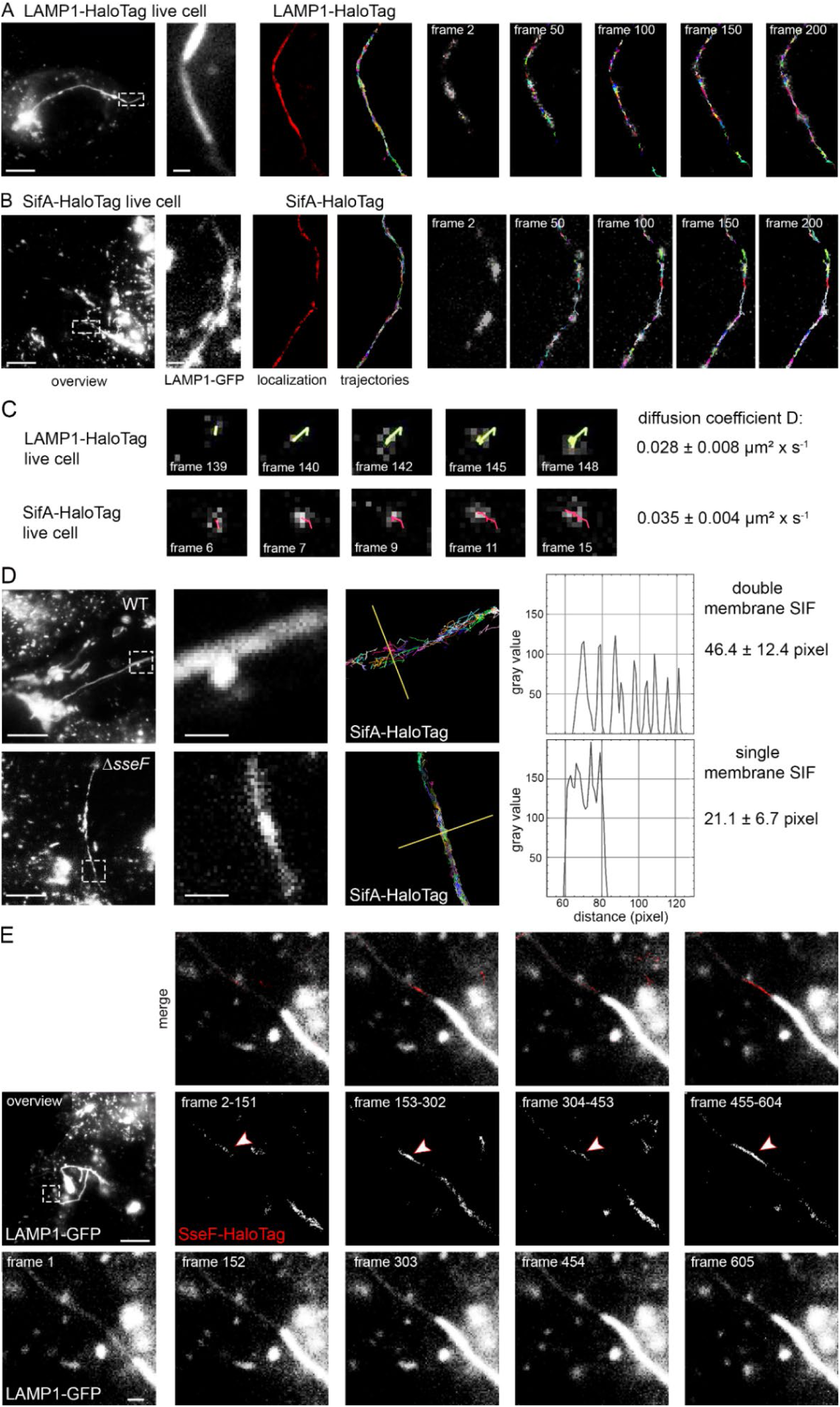
Single molecule localization and tracking of STM SPI2-T3SS effector proteins on single-membrane SIF. HeLa cells stably expressing LAMP1-GFP were infected with STM *sseF, sifA* mutant strains expressing *sifA*::HaloTag::HA and labeled with HTL-TMR as described above. For visualization of LAMP1-HaloTag, the cells were transfected with LAMP1::HaloTag::HA one day before infection, and infected with STM *sseF* mutant strain. **A, B)** Representative SRM images acquired using 15% laser power at the focal plane, rendered from single molecule localization and tracking within 200 consecutive frames. Selected frames (frame rate: 32 frames per second) of the TMR signal, localization and tracking are presented, (also showing elapsed trajectories). The sequences of 200 frames of SifA-HaloTag and LAMP1-HaloTag are shown in **Movie 7** and **Movie 8**. **C)** Selected frames of trajectories from a single molecule. Using at least 2,800 pooled trajectories for proteins in at least 20 infected cells in three biological replicates recorded under the same conditions, the diffusion coefficient D was calculated applying the Jaqaman algorithm. The indicated error is the calculated error of the resulting slope (with 95% confidence bounds). **D)** Intensity profile analysis of SifA-HaloTag trajectories on sm and dm SIF. The intensity profiles of trajectories tracked on SIF were analyzed using the Fiji plot profile tool. SIF from various infected cells were processed and resulting pixel range of the profile was determined. **E)** HeLa cells stably expressing LAMP1-GFP were infected with STM *sseF* mutant strain expressing *sseF*::HaloTag::HA and labeled with HTL-TMR as described above. The transition of leading to trailing SIF was imaged with 488 nm laser excitation for 1 frame (frame rate: 32 frames per second) following 561 nm laser excitation for 150 frames in 4 cycles. Shown are representative SRM images acquired using 15% laser power at the focal plane, rendered from SML within each of the 150 consecutive frames. High local concentration of effector protein on leading SIF is indicated by arrowheads. The sequences of 5 frames of LAMP1-GFP are shown in. Scale bars: 10 and 1 µm in overviews and details, respectively.

Taken together, these single molecule analyses demonstrate that SPI2-T3SS effector proteins are highly dynamic on SIF. PipB2 showed distinct higher mobility in comparison to host membrane integral protein LAMP1. The mobility of SifA and LAMP1 was reduced on sm SIF in comparison to dm SIF.

### SPI2-T3SS effectors accumulate on leading SIF during transition to trailing SIF

After succeeding in imaging effectors on sm SIF, we analyzed the transition of leading to trailing SIF. Thinner leading SIF consist of single-membrane tubules and the connected trailing SIF consist of double-membrane structures, i.e. fully developed SIF. It was proposed that this transition is facilitated by a lateral extension of membranes of leading SIF, engulfment of host cell cytosol and cytoskeletal filaments, and finally membrane fusion to form double-membrane trailing SIF (Krieger et al., 2014). To image the transition from leading to trailing SIF, translocated SseF-HaloTag was analyzed by LCI at 6 h p.i. In **Fig. 5E** and **Movie 9**, a thin SIF with a weak GFP signal was imaged, and a wider trailing SIF with strong GFP signal was developing alongside the thinner structure. By collecting and localizing all signals of SseF-HaloTag between the frames of the growing LAMP1-positive SIF tubule, increased concentration of effector proteins on the leading SIF before transition to trailing SIF became apparent. This was also shown for SifA-HaloTag and PipB2-HaloTag (**Fig. S 4AB, Movie 10, Movie 11**). These effector proteins were also localized at leading SIF when transforming to trailing SIF, however the local concentration was less pronounced.

These findings indicate that SPI2-T3SS effector proteins are already present on leading SIF, and in particular SseF appears to be involved in the transformation to trailing dm SIF, as an accumulation of effector protein can be detected directly before transition.

### Effector proteins target endosomal vesicles in the early phase of infection

The presence and concentration of effector proteins on leading SIF suggest a delivery mechanism of effectors to the tips of growing sm SIF tubules. To address the question how SPI2-T3SS effector proteins reach their subcellular destination in an infected host cell, we applied LCI by confocal laser-scanning microscopy (CLSM) of infected HeLa LAMP1-GFP cells at 4 h p.i. In the early stage of infection, the SCV is already formed, while SIF biogenesis initiates. We found that after labeling of effector proteins, also bacteria were heavily stained, indicating large amounts of effector proteins stored in bacteria. By monitoring the HaloTag-fused PipB2, SseF, SseJ and SteC, localization of effector proteins in a punctate, vesicle-like manner was observed in infected cells. These structures showed most frequently colocalization with LAMP1-GFP signal, but also labeled endomembrane compartments lacking the late endosomal marker were observed (**Fig. S 5**).

SifA-HaloTag was not detected decorating vesicles in the early stage of infection. The low level of SifA-HaloTag translocation could hamper visualization. These findings are in line with the observation made by SRM localization with SifA-HaloTag showing lowest effector concentration, while PipB2-HaloTag showed the highest labeling intensity on SIF tubules. Accordingly, vesicles marked with PipB2-HaloTag could be easily imaged in the early stage of infection, and PipB2 protein was chosen for further analyses. We set out to determine different phenotypes of PipB2-HaloTag localization and therefore studied 100 cells containing PipB2-HaloTag-positive vesicles. Of note, at 4 h p.i. effector-positive vesicles were found in a subset of infected cells. We conclude that due to heterogeneity in SPI2 induction, infected cells with bacteria with low levels of effector secretion showed no detectable HaloTag signal. In line with this observation, PipB2-HaloTag fluorescence intensity on SIF at 16 h p.i. also varied between infected cells (**Fig. S 6**). In the early phase of infection, distinct phenotypes of PipB2-HaloTag localization can be distinguished for effector-positive vesicular structures. Moreover, the infected cells either showed no SIT, or already developing first SIT structures. The tubular structures were either LAMP1-GFP-positive, or lacking the endosomal marker, and in one population of cells these structures had already acquired PipB2-HaloTag, and others were still lacking the effector (**Fig. 6A**). At 4 h p.i., in total 62% of the infected cells still did not show SIF formation, yet were positive for vesicles decorated with PipB2-HaloTag. All other cells also displayed vesicles positive for PipB2-HaloTag and already formed SIT (**Fig. 6B**). To test the characteristics of effector-decorated vesicles in infected cells, we performed tracking analyses of vesicles. Single LAMP1-GFP-positive or PipB2-HaloTag-positive vesicles were tracked in 3D in infected cells. In co-motion tracking analyses, we observed that vesicles positive for LAMP1-GFP and PipB2-HaloTag were tracked in parallel, and the patterns of movement were identical (**Fig. 6AB, Movie 12**). When studying individual vesicle tracks, as control conditions LAMP1-GFP-decorated vesicles in non-infected cells, either nocodazole-treated or non-treated were tracked (**Fig. S 6, Movie 13, Movie 14**).

**Fig. 6.**
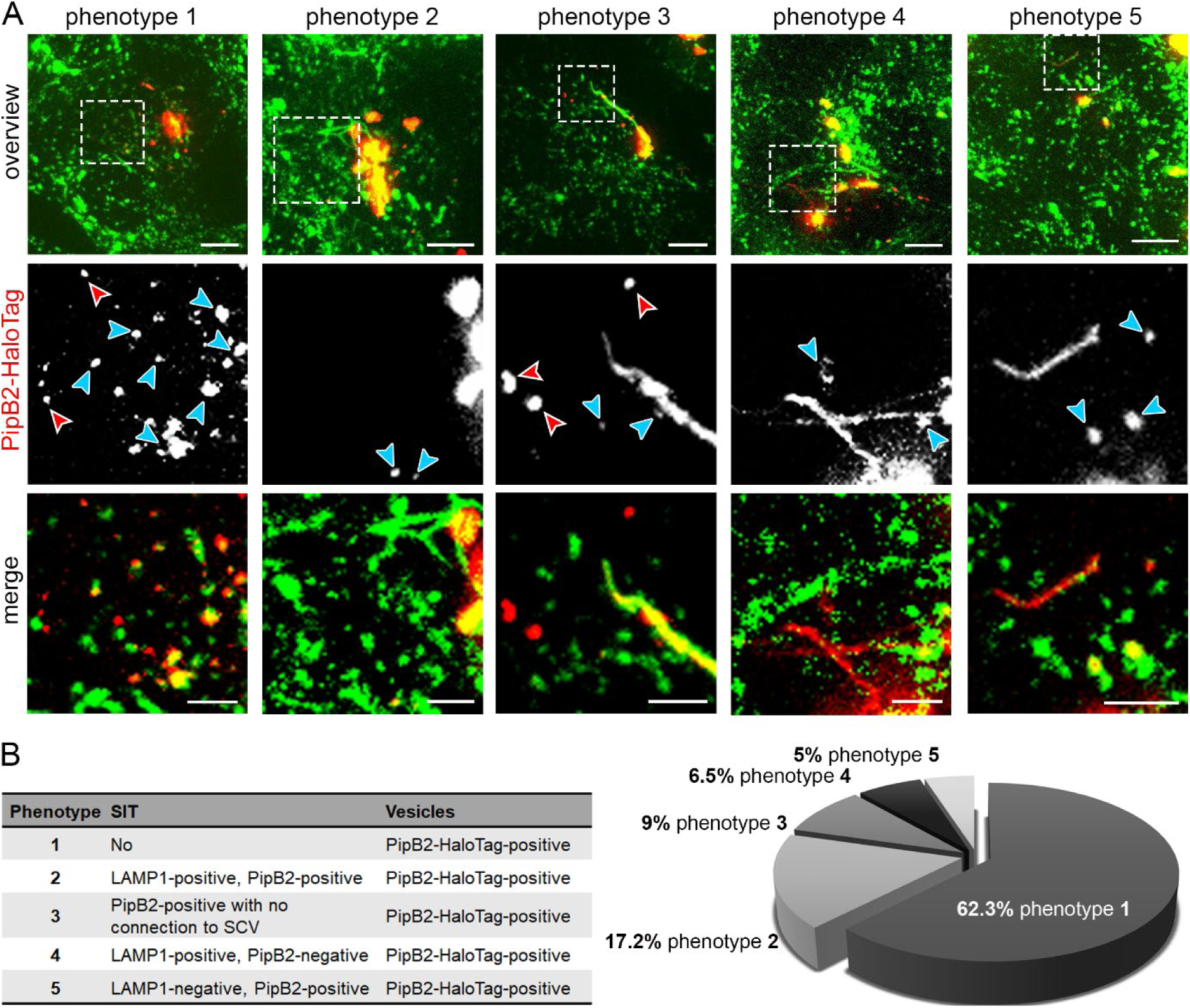
Distribution of PipB2-HaloTag in the early phase of infection. HeLa LAMP1-GFP cells were infected with STM *pipB2* mutant strain expressing *pipB2*::HaloTag::HA. **A)** LCI was performed directly after cells were stained with 1 µM HTL-TMR at 3.5 h p.i. for 30 min. For at least 100 infected cells with PipB2-HaloTag-positive vesicles, the phenotypes of PipB2-HaloTag localization on vesicles and SIT were determined. Blue arrowheads indicate vesicles double-positive for LAMP1-GFP and HTL-TMR-labeled PipB2-HaloTag. Red arrowheads indicate vesicles negative for LAMP1-GFP and positive for PipB2-HaloTag. Scale bars: 10 and 2 µm in overviews and details, respectively. **B)** Quantification of distinct PipB2-HaloTag distributions in infected HeLa LAMP1-GFP cells.

We quantified the mean track displacement length (MTDL) and the mean track speed (MTS) of pooled trajectories. In infected cells, PipB2-HaloTag-marked and LAMP1-GFP-marked vesicles did not differ in MTDL and MTS and therefore showed normal characteristics of vesicle movement. In cells treated with nocodazole, both values were significantly decreased due to inhibition of vesicle trafficking after microtubule disruption. Interestingly, late endosomal vesicles tracked in non-infected and non-treated cells showed a more rapid movement, and MTDL and MTS values were significantly increased (**Fig. 7C**).

**Fig. 7.**
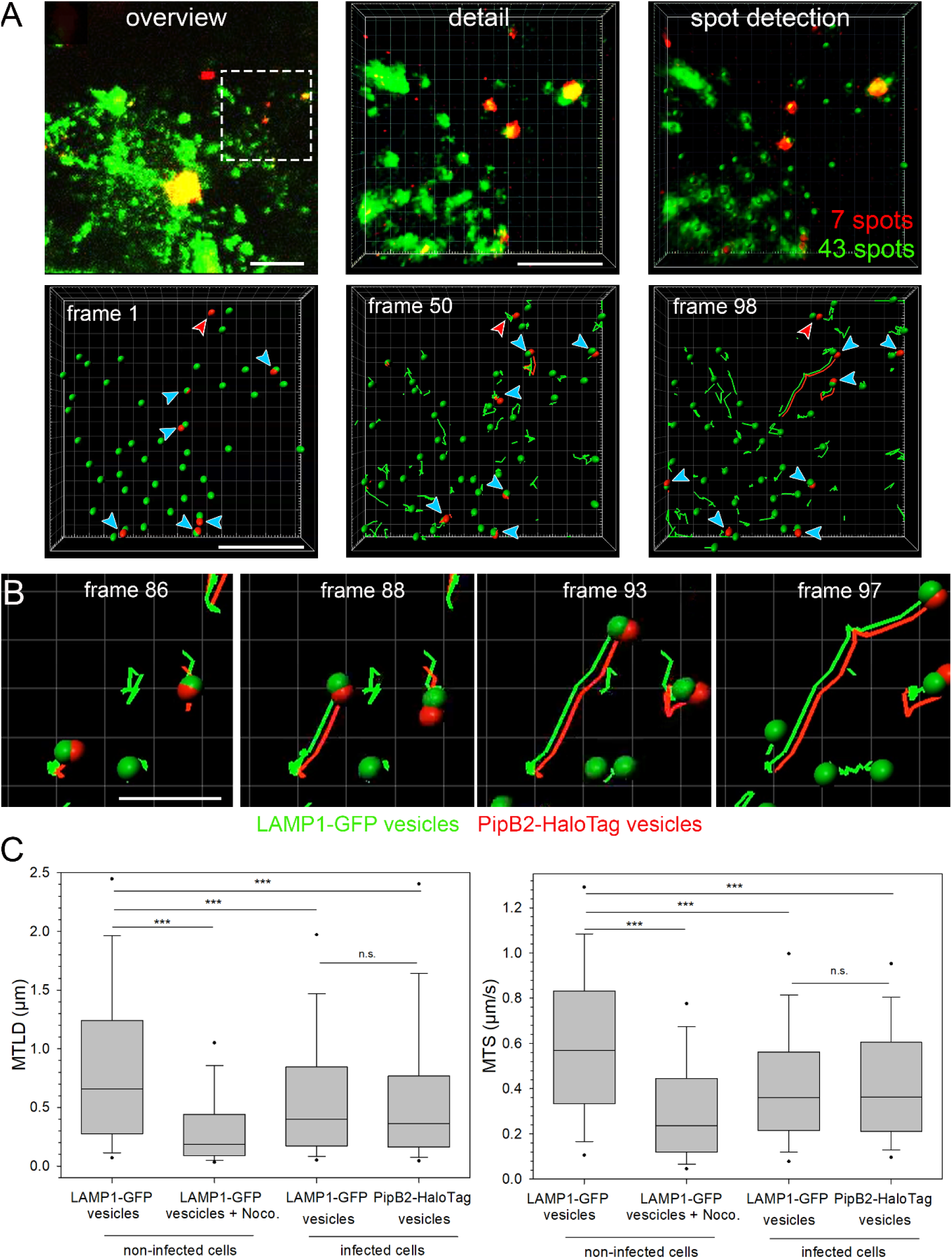
Tracking of vesicles positive for LAMP1-GFP and PipB2-HaloTag. HeLa LAMP1-GFP cells were either not treated, treated with nocodazole to inhibit vesicle movement, or infected with STM Δ*pipB2* strain expressing *pipB2*::HaloTag::HA. **A)** An infected HeLa LAMP1-GFP cell with LAMP1-GFP (green) and PipB2-HaloTag-TMR (red) was imaged for 200 frames (0.39 frames/sec) by SDM in dual camera streaming mode. Vesicle tracking analysis was done with the Imaris spot detection tool and co-motion analysis is shown at different time points (**Movie 12**). Blue arrowheads indicate vesicles positive for LAMP1-GFP and effector protein fused to HaloTag and labeled with TMR. Red arrowheads indicate vesicles negative for LAMP1-GFP and positive for effector protein. **B)** Trajectories of single vesicles labeled with LAMP-GFP and PipB2- HaloTag. Scale bars: 10 and 5 µm in overviews and details, respectively in **A**, 2 µm in **B**. **C)** Quantification of at least 858 trajectories from five individual cells per condition. Box plot analysis of mean track displacement length (MTDL) and mean track speed (MTS) of vesicles under various conditions. Statistical analyses were performed by Rank Sum test and significances are indicated as follows: n.s., not significant, ***, *p* < 0.001.

These data demonstrate that SPI2-T3SS effectors are recruited to endomembrane compartments, and as analyzed in detail for PipB2-HaloTag, in infected cells in the early phase of infection PipB2-HaloTag-positive vesicles behave similar to LAMP1-positive vesicles.

### PipB2-HaloTag-positive vesicles continuously integrate into the SIF network

After establishing that SPI2-T3SS effector proteins are recruited to vesicles in the early infection during SIF biogenesis, we hypothesized that delivery to SCV and SIF tubules may occur by fusion of endosomal vesicles with membrane-integrated effector proteins. We set out to study PipB2-HaloTag localization from early to late stage of infection and applied long-term LCI of infected cells. Over time, we observed a reduction of effector-decorated vesicles and extension of effector-positive SIF network (**Fig. 8A**, **Movie 15**). Rendering the PipB2-HaloTag-labeled endomembranes, 29 effector-positive vesicle-like objects were detected 5 h p.i. while only six objects remained at 12 h p.i. Concurrent with the decreasing number of PipB2-HaloTag-decorated objects, PipB2-HaloTag-positive SIF developed (**Fig. 8C**), suggesting that effector-positive SIF membranes emerge from vesicles. We speculate that STM mutant strains without the ability to induce tubulation of SIF but still possessing a functional SCV should accumulate effector-positive vesicles over time. We used STM deficient in *sifA* and *sseJ,* previously reported to maintain SCV membrane but lacking SIF formation (Ruiz-Albert *et al*., 2002). In infected HeLa LAMP1-GFP cells at 16 h p.i. an accumulation of PipB2-HaloTag-positive vesicles was observed, as well as lack of SIF network formation (**Fig. S 5**). These findings indicate that STM SPI2-T3SS effector proteins are integrated into the SCV-SIF continuum via fusion of effector–positive endomembrane compartments.

**Fig. 8.**
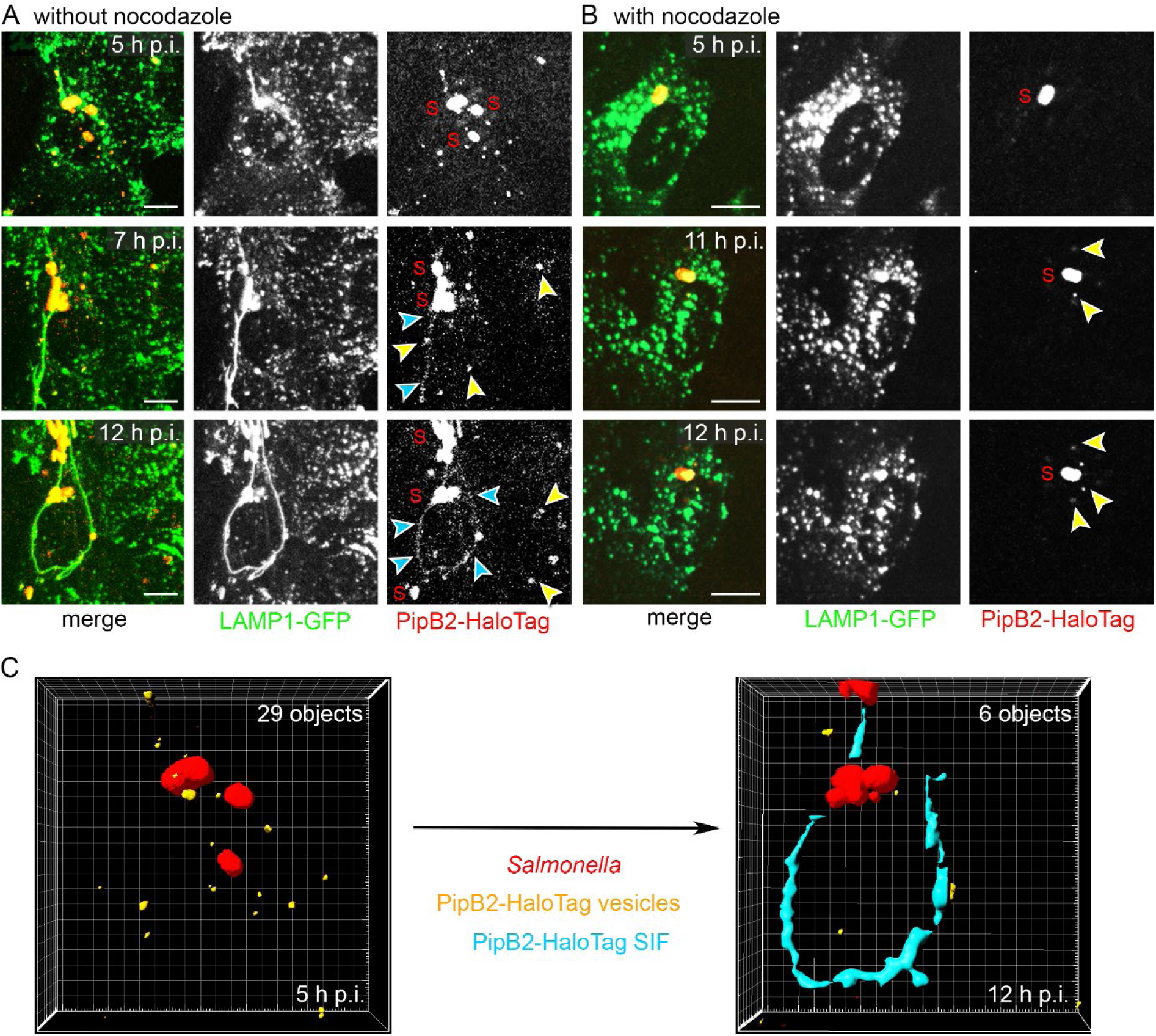
Conversion of vesicular to tubular distribution of translocated effector proteins. HeLa LAMP1-GFP cells were infected with STM Δ*pipB2* strain expressing *pipB2*::HaloTag::HA. Cells were either not treated (**A**), or treated with nocodazole (5 µg x ml ^-1^) 2 h p.i. (**B**). The inhibitor was removed after HaloTag staining and cells were washed twice. LCI was performed using SDM directly after cells were stained with 1 µM HTL-TMR for 30 min. The cells were imaged over a period of 8 h every 30 min (**Movie 15, Movie 16**). Representative STM are labelled (S), and PipB2- HaloTag-positive vesicles or SIF are indicated by yellow or blue arrowheads, respectively. Scale bars: 5 µm. **C)** The Imaris surface analysis tool was used to determine in an infected cell at 5 h and 12 h p.i. the amounts of either vesicles (orange), or SIF tubular structures (blue) positive for PipB2-HaloTag.

### Effector recruitment to endomembrane compartments is dependent on vesicle movement

To investigate if presence of effector-decorated vesicles is dependent on dynamics of endosomal compartments, we monitored localization of PipB2-HaloTag over time in infected cells after inhibition of vesicle trafficking. For this purpose, infected cells were treated with nocodazole 2 h p.i., a time point where STM resides in SCV and activates expression of SPI2-T3SS genes (Hapfelmeier et al., 2005). Addition of nocodazole abrogates vesicle movement due to interference with microtubule integrity. We did not observe PipB2-HaloTag-positive vesicles in nocodazole-treated cells, implicating that vesicle movement is required for effector recruitment **Fig. 8B**, **Movie 16**).

To address reversibility of inhibition, we imaged infected cells for 8 h after removing nocodazole by washing cells twice at 4 h p.i. A slow initiation of vesicle movement was monitored over time. Starting 11 h p.i., the first PipB2-HaloTag-decorated LAMP1-GFP-positive vesicles were imaged (**Fig. 8B**). Association of effector proteins with endosomal compartments was dependent on vesicle movement on microtubules, but did not require SIF formation.

## Discussion

In this study, we demonstrated that SPI2-T3SS effector proteins, as well as the host cell protein LAMP1 can be tracked, on single molecule level, on SIF and show a rapid, bidirectional movement. The ability of integral membrane proteins to rapidly interchange in SIF membranes has already been demonstrated by FRAP experiments and that revealed recovery of LAMP1 after photobleaching on distal SIF (Liss et al., 2017). Using the split-GFP approach, PipB2 recovery on tubules was detected upon photobleaching, indicating a rapid distribution of PipB2 (Van Engelenburg & Palmer, 2010). The diffusion coefficients (DC) are similar to values determined for mitochondrial proteins and for cytokine receptors on the plasma membrane (Appelhans & Busch, 2017; Richter et al., 2017).

Interestingly, the mobility of effectors differs with PipB2 being the most mobile effector protein. Mobility of membrane proteins depends on membrane integration, membrane composition and interaction with other proteins. The association of PipB2 to lipid rafts has been demonstrated (Knodler et al., 2003). These dense membrane patches would presume an impaired mobility for membrane integral proteins. Contrary, it is established that raft association is not the dominant factor in determining the overall mobility of a particular protein as different lipid raft associated proteins showed diverse DC (Kenworthy et al., 2004). Besides this, our experiments did not reveal a specific localization of PipB2 to distinct regions of SIF membranes.

Correlation of different forms of protein association with membranes to the corresponding diffusion mobility revealed that DC of prenylated proteins were higher than DC of transmembrane proteins (Kenworthy et al., 2004). We did not observe such a correlation for STM effector proteins, with SifA being the only prenylated effector protein tested (Reinicke et al., 2005), and propose that mobility of effectors might be influenced to a large extend by their interaction with host proteins. SseF interacts with STM effector SseG and in combination with the Golgi network-associated protein ACBD3, forming a complex tethering the SCV to the Golgi (Yu et al., 2016). SifA has various interaction partners inside the host cell, such as PLEKHM2 (also called SKIP) and PLEKHM1 (Boucrot et al., 2005; McEwan et al., 2015). The SifA complex is able to activate kinesin-1 upon binding, resulting in budding and anterograde tubulation of SIF membranes along microtubules (Dumont et al., 2010; Schroeder et al., 2011; Yip et al., 2016). PipB2 is responsible of the recruitment of auto-inhibited kinesin-1 to the vacuole membrane (Henry et al., 2006). The distinct DC of effector proteins SifA, SseF and PipB2 likely indicates the form of interaction with cognate host cell molecules. Both, SifA and SseF, take part in tethering SCV and SIF to host structures, possibly resulting in reduced mobility. Hence, the molecular mechanisms behind effector mobility on SIF membrane remain to be elucidated.

We demonstrate here that effector motilities are affected by SIF architecture. The reduced DC of STM effector protein SifA, as well as host protein LAMP1 reveals slower movement on sm SIF. This indicates an overall impediment of mobility of membrane proteins in the membrane of sm SIF. The smaller diameter of sm SIF compared to dm SIF leads to higher membrane curvature of sm SIF, and this may affect motility of membrane proteins. Furthermore, host cell proteins sensing membrane curvature will be recruited differentially to sm SIF and dm SIF depending on their curvature, and these proteins could have distinct effects on motilities of the proteins analyzed.

When monitoring growing SIF, we found thinner leading SIF at the extending tips, followed by development of thicker trailing SIF. Prior ultrastructural analysis revealed that leading SIF are single-membrane tubules, while trailing SIF comprise double-membrane structure, representing the fully developed SIF. It was proposed that effector protein SseF together with its interaction partner SseG are involved in conversion of sm SIF to dm SIF (Krieger et al., 2014). Here we present, to our knowledge first, microscopy-based evidence that effector protein SseF is present on the membranes of leading sm SIF and that the effector accumulates before engulfment to trailing dm SIF. As this is also true for SifA and PipB2, more effector proteins might be involved in the process of sm SIF to dm SIF conversion.

The concentration of effectors on the tips of growing SIF led to the question how STM effectors are able to accumulate and how they are recruited after secretion into the host cytosol. Especially SseF which is characterized by large hydrophobic domains (Abrahams et al., 2006), could be mistargeted to cytoplasmic membranes or be prone to aggregation in host cytosol. To test models for the delivery of effector proteins to SCV-SIF continuum introduced in **Fig. 3**, we performed LCI with labeled effectors in the early infection. We found that various SPI2-T3SS effectors decorate host endomembranes in infected cells. The association of PipB2 with vesicles in the periphery of infected host cells has already been demonstrated (Knodler et al., 2003). Over time, the presence of PipB2-positive vesicles declined, corresponding with the formation of PipB2-positive SIF. These findings strengthen the hypothesis of delivery of effector proteins to SCV and SIF via fusion of effector-containing endomembranes as depicted in models B and C of **Fig. 3**.

The targeting of bacterial effector proteins to phagosomes and various organelles in infected host cells has been widely demonstrated (Popa et al., 2016), but only few studies observed effector protein presence on host endomembranes and connected a trafficking mechanism of effector proteins to these phenomena. The *Shigella* effector protein IpgB1 was found to localize to endocytic vesicles in mammalian cells expressing eGFP-tagged IpgB1. Vesicles decorated with IpgB1 were found to be functional in host cell trafficking and a relocation of effector proteins from endocytic vesicles to membrane ruffles produced by *Shigella* was observed, indicating a delivery of effector via vesicle fusion (Weigele et al., 2017). Another study uses the split GFP approach to monitor the delivery of effector AvrB by *Pseudomonas* via the T3SS to infected plant cells. In infected host cells, AvrB localizes to the plasma membrane, but at different time points after inoculation the localization changed to unknown vesicles, suggesting a potential trafficking of AvrB on vesicles (Park et al., 2017). Relocation of bacterial effector proteins on host endocytic vesicles to the required site of action in infected host cells might represent a universal delivery mechanism.

To further address the question how effector proteins reach their subcellular destination on host endomembranes, we analyzed effector protein localization in infected cells with inhibited vesicle movement. We found that endomembranes did not acquire PipB2 when vesicle movement was inhibited. This finding supports an insertion route of effector proteins to membranes that requires vesicle movement. In this case, effector proteins which do not require a modification by host proteins are inserted into vesicle membranes in close proximity to the T3SS, either by host chaperones, or solely determined by the sequence of membrane integral effectors (**Fig. 3**, model C).

However, these hypotheses can only be applied to effector proteins without requirement for host cell modification. SPI2-T3SS effector proteins can gain membrane association either by defined trans-membrane domains within their sequence, or by host cell modifications. The effector proteins SseF and SseG are examples for the former, as these proteins contain identified trans-membrane domains (Hansen-Wester et al., 2002; Hensel et al., 1998; Krampen et al., 2018). Various other SPI2-T3SS effector proteins acquire membrane association by host cell modifications. These modifications comprise different mechanisms to increase the overall hydrophobicity of effector proteins. SifA is targeted by host modifications, resulting in S-prenylation of the effector protein (Reinicke et al., 2005). Additionally, the effector proteins SspH2 and SseI of STM have been shown to be palmitoylated to gain membrane association (Hicks et al., 2011). As the mentioned models do not include effectors requiring post-translational modification by the host, other mechanisms must contribute to effector relocation (**Fig. 2**, model D). Accordingly, we did not detect SifA on host endomembranes in infected cells.

Our data lead to various new questions. i) During the intracellular life of STM, host endosomal compartments are gradually depleted due to fusion to SCV-SIF continuum (Rajashekar et al., 2008). Does this terminate the delivery of effector proteins and end intracellular proliferation? ii) By which mechanism are post-translationally-modified effector proteins relocated? iii) Are endomembranes specifically targeted to the T3SS, or is insertion simply dependent on proximity? Further LCI and single molecule-based analyses of STM effectors will likely contribute to answer these questions.

## Acknowledgements

This work was supported by the DFG through grants HE 1964/18-2 and SFB 944 project Z to M.H. We like to thank Jacob Piehler (Div. Biophysics), Christian Richter (Div. Biophysics) and Rainer Kurre (iBiOs) for continuous support and fruitful discussions.

## Materials and Methods

### Bacterial strains and culture

Infection experiments were performed using *Salmonella enterica* serovar Typhimurium (STM) NCTC 12023 strain as WT and isogenic mutant strains (Table 1). Mutagenesis was carried out as described elsewhere (Popp et al., 2015). In short, strains were constructed using λ Red-mediated mutagenesis and resistance cassette was removed using FLP-mediated recombination. Mutant strains deficient in effector genes harboring plasmids encoding the corresponding effector fused to HaloTag (Table 2) were used for microscopic analysis. Plasmids were constructed as described previously (Göser et al., 2019) using oligonucleotides listed in Table S 1. Bacterial strains were cultured in Luria-Bertani broth (LB) containing 50 µg x ml^-1^ Carbenicillin.

**Table 1.**
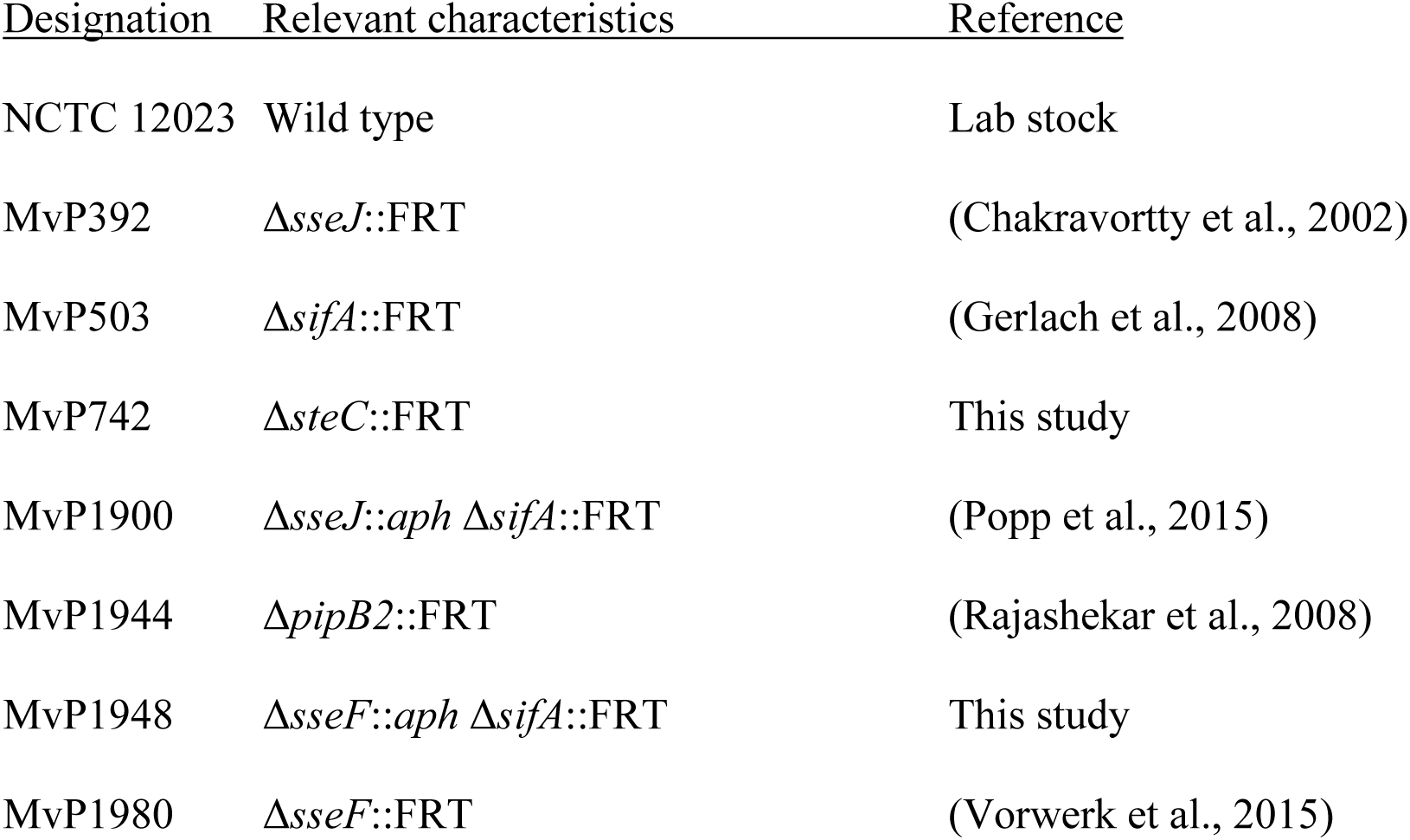
*Salmonella enterica* serovar Typhimurium strains used in this study.

**Table 2.** Plasmids used in this study.

| 706 | <u>Designation</u> | <u>Relevant genotype</u> | <u>Source/Reference</u> |
| --- | --- | --- | --- |
| 707 | p2095 | $P_{sseA::sscB} sseF::M45$ | (Kuhle et al., 2004) |
| 708 | p2129 | $P_{sseJ} sseJ::M45$ | (Hansen-Wester et al., 2002) |
| 709 | p2621 | $P_{pipB2} pipB2::M45$ | (Knodler et al., 2003) |
| 710 | p2643 | $P_{sseA::sscB} sseF::HA$ | (Kuhle et al., 2004) |
| 711 | p3991 | LAMP1::HaloTag::GFP | (Liss et al., 2017) |
| 712 | p4305 | $P_{sifA} sifA::L16::HaloTag::HA$ | (Göser et al., 2019) |
| 713 | p4118 | $P_{sseA} sscB sseF::L16::HaloTag::HA$ | (Göser et al., 2019) |
| 714 | p4286 | $P_{sseJ} sseJ::L16::HaloTag::HA$ | (Göser et al., 2019) |
| 715 | p4295 | P<sub>*pipB2*</sub> *pipB2*::L16::HaloTag::HA | (Göser et al., 2019) |
| 716 | p5059 | P<sub>*steC*</sub> *steC*::L16::HaloTag::HA | This study |
| 717 | p5065 | P<sub>*sseA*</sub>::*sscB sseF*::3HA | This study |
| 718 |  |  |  |

For generation of a plasmid encoding triple HA-tagged SseF, p2643 (*sscB sseF*::HA) was used and *sseF*::HA on p2643 was replaced by *sseF*::3HA using Gibson assembly GA. Primers for generation of vector fragment, check primers and sequence of synthetic *sseF*::3xHA (gBlocks, IDT) are listed in Table S 1.

### Culture of eukaryotic cells

The non-polarized epithelial HeLa cell line (ATCC no. CCL-2) stably transfected with LAMP1-GFP (Krieger et al., 2014) was cultured in Dulbeccós modified Eaglés medium (DMEM) containing 4.5 g x l^-1^, glucose, 4 mM stable glutamine and sodium pyruvate (Biochrome) and supplemented with 10% inactivated fetal calf serum (Sigma-Aldrich). Cells were maintained in a cell culture incubator (37 °C, 5% CO_2_).

### Host cell infection

HeLa LAMP1-GFP cells were seeded in surface-treated 8-well plates (ibidi) or on 24 mm diameter coverslips (VWR) in 6-well plates (Faust). For infection experiments cells were grown to 80% confluence (8 well: ∼ 8 x 10^4^, 6 well: ∼ 6 x 10^5^). STM strains were grown over night in LB and subcultured 1:31 in fresh LB for 3.5 h. Cells were infected at MOI 50 or 75 for 25 min, washed thrice with phosphate-buffered saline (PBS) and incubated with DMEM containing 100 µg x ml^-1^ gentamicin (Applichem) to kill non-internalized bacteria. Subsequently, medium was replaced by DMEM containing 10 µg x ml^-1^ gentamicin for the rest of the experiment.

### Pulse-chase with gold nanoparticles

Pulse-chase labeling of the endosomal system with rhodamine conjugated gold nanoparticles was performed as previously described (Zhang & Hensel, 2013).

### Correlative light and electron microscopy (CLEM)

CLEM of HeLa LAMP1-GFP cells infected by STM WT was performed as previously described (Krieger et al., 2014). Briefly, HeLa LAMP1-GFP cells were grown on MatTek dishes with a gridded coverslip and infected with the respective STM strains at MOI 75. Cells were fixed with 2% paraformaldehyde (PFA) and 0.2% glutaraldehyde (GA) in 0.2 M HEPES for 30 min. prior to LM. After rinsing the cells for three times with 0.2 M HEPES buffer, unreacted aldehydes were blocked by incubation with 50 mM glycine in buffer for 15 min, followed by rinses in buffer. CLSM was performed and ROIs were chosen. Afterwards cells were fixed with 2.5% glutaraldehyde and 5 mM CaCl_2_ in 0.2 M HEPES in preparation for TEM. Further steps including post-fixation, dehydration, sectioning and imaging in the TEM were conducted as previously described (Krieger et al., 2014).

### Live-cell super-resolution localization and tracking microscopy

Localization and tracking of single molecules were done as previously described (Göser *et al*., 2019). Briefly, infected HeLa LAMP1-GFP cells were labeled with the HaloTag ligand coupled to tetramethylrhodamine (HTL-TMR, Promega) in a concentration of 20 nM for 15 min at 37 °C. After ten washing steps, the cells were imaged with an imaging medium consisting of Minimal Essential Medium (MEM) with Earle’s salts, without NaHCO_3_, without L-Glutamine, without phenol red (Biochrom) and supplemented with 30 mM HEPES (Sigma-Aldrich), pH 7.4. TIRF microscopy was performed using an inverted microscope Olympus IX-81, equipped with an incubation chamber maintaining 37 °C and humidity, a motorized 4-line TIRF condenser, a 150 x objective (UAPON 150x TIRF, NA 1.45), a TIRF quadband polychroic mirror (zt405/488/561/640rpc), a 488 nm laser (150 mW, Olympus), and a 561 nm laser (150 mW, Olympus). Localization as well as tracking of single molecules were carried out with the help of a self-written user interface in MatLab 2013a (Appelhans et al., 2018; Barlag et al., 2016; Hess et al., 2006; Jaqaman et al., 2008; Serge et al., 2008; Wilmes et al., 2012).

### Live cell fluorescence microscopy

Infected HeLa LAMP1-GFP cells were labeled with HTL-TMR with a final concentration of 1 µM for 30 min directly before LCI. Cells were washed thrice with PBS and the media was replaced with imaging media as described above. For inhibition of vesicle movement cells were incubated with 5 µg x ml^-1^ nocodazole (Sigma-Aldrich) 2 h p.i. Directly before LCI, cells were washed twice to remove nocodazole and labeled as described above. Imaging was done using either the confocal laser scanning microscope Leica SP5 (Leica) equipped with an incubation chamber, a 100 x objective (HCX PL APO CS, NA 1.4-0.7) and a polychroic mirror (TD 488/543/633) or the Cell Observer Spinning Disk microscope (SDM, Zeiss) equipped with a Yokogawa Spinning Disc Unit CSU-X1a 5000, an incubation chamber, 63 x objective (α-Plan-Apochromat, NA 1.4), two EMCCD cameras (Evolve, Photometrics) and the filter combination GFP with BP 525/50 and RFP with BP 593/46 for dual cam imaging. Images were acquired and processed using the corresponding software LAS AF (Leica) and ZEN (Zeiss).

### Image analysis by Imaris

Vesicle tracking and surface analysis were analyzed by Imaris 9.2.1 software package (Bitplane). Surface analysis was done to determine the amounts of effector-positive vesicles inside an infected cell. By the surface tool, data acquired by live cell imaging were analyzed in the red (PipB2-HaloTag-TMR) channel. Vesicle volume was adjusted using auto-threshold and smoothing of 0.1, and SIF volume was adjusted using auto-threshold and smoothing of 0.3. For vesicle tracking the spot tool was used. Spot detection was performed with the following parameters XY diameter: 0.75 µm, active model PSF elongation: 1.5 µm and background subtraction. For tracking the autoregressive motion algorithm was chosen with a maximum gap size of 3.

### Image analysis by Fiji

The Fiji package (Schindelin et al., 2012) was used to determine intensity profiles of accumulated SifA-HaloTag trajectories on SIF. Using the line tool and subsequently the plot profile tool, the intensity profile of the trajectories labeling the SIF was calculated and the relative distances in pixel were compared.

## Suppl. Materials and Methods

### Western blot analysis

STM strains were cultured overnight (o/n) in PCN, 1 mM P_i_, pH 7.4 medium, diluted 1:31 in fresh PCN, 0.4 mM P_i_, pH 5.8 medium and subcultured for 6 h. Subsequently, the optical density at 600 nm (OD_600_) was measured and 300 µl of bacterial suspension was transferred to a 1.5 ml tube. Bacteria were pelleted by centrifugation (22,000 x g, 2 min, 4 °C), resuspended in 1 x SDS sample buffer adjusted to 1 unit of OD_600_ per 100 µl and lysed by incubation at 100 °C for 5 min. Samples of 10 µl were subjected SDS-PAGE at 150 V for 75 min. Semi-dry blotting onto 0.45 µm nitrocellulose membranes was performed at 10 V constant for 45 min. Blocking of membranes was performed with 5% milk powder in TBS/T (0.1% Tween 20 in TBS) for 60 min at RT. Subsequently, primary antibody against HA tag was incubated (1:10,000 in TBS/T) for 1 h at RT, followed by incubation of secondary HRP-coupled antibody (1:10,000 in TBS/T) for 1 h at RT. Antibodies used in this study are listed in Table S 2. Between incubation with antibodies, membranes were washed thrice with TBS/T for at least 10 min each. Signals were recording using the ChemiDoc system from BioRad and its corresponding software ImageLab.

### Cryo sample preparation for Tokuyasu immunogold labeling and TEM

Two days prior to infection HeLa LAMP1-GFP cells (1.5 x 10^6^) were seeded into a 60.1 cm^2^ Petri dish. cells were pre-fixed at 8 h p.i. for 10 min. with pre-warmed double-concentrated fixative (4% (w/v) PFA, 0.2% (v/v) GA) in 0.1 M PHEM buffer which was added to the culture dish 1:1 mixed with the culture medium. Subsequently, fixative was replaced by fresh fixative (2% (w/v) PFA, 0.1% (v/v) GA) and cells were fixed for 2 h at RT and stored o/n in 1% formaldehyde (w/v). Next, cells were washed twice with 0.1% glycine in 0.1 M PHEM, several times with 0.1 M PHEM buffer, scraped in PHEM containing 1% (w/v) gelatin from the culture plates and were pelleted by centrifugation (300 x g, 3 min). The cell pellet was infiltrated at 37 °C stepwise in 2% (w/v), 5% (v/w) and finally in 10% (w/v) gelatin in 0.1 M PHEM buffer. After gelation at 4 °C, 1 mm^3^ cubes were dissected and infiltrated in 2.3 M sucrose o/n at 4 °C in rotating vials. Gelatin cubes were mounted on specific aluminum specimen holders and plunge-frozen in liquid nitrogen. Specimen holders were placed into the cryo-chamber of a cryo-ultramicrotome UC7 (Leica Microsystems, Wetzlar), precooled to -110 °C and trimmed to suitable block size. Ultrathin sections of 60 nm were cut at -110 °C with a dry cryo-immuno diamond knife (Diatome, Switzerland). Ribbons of sections were picked up with a wire loop filled with a 1+1 mixture of 1% (w/v) methyl cellulose and 2.3 M sucrose in PHEM buffer. Sections were thawed on the pick-up solution and transferred downwards to Formvar carbon-coated 100-mesh copper grids.

For immunolabeling sections were placed 30 min on 37 °C warm water to diffuse pick-up solution and gelatin. Subsequently, grids were rinsed over a series of droplets: washed in 0.1% glycine in PBS, blocked 3 min in 1% BSA in PBS, incubated 60 min. in primary antibody diluted in 1% BSA, 0.2% fish skin gelatin in PBS, washed in 0.1% BSA in PBS, incubated 30 min in bridging antibody, diluted in 1% BSA, 0.2% fish skin gelatin in PBS, washed in 0.1% BSA in PBS, incubated 20 min in 10 nm protein A-gold diluted in 1% BSA in PBS, washed in PBS, fixed 5 min in 1% (v/v) glutaraldehyde in PBS and washed in distilled water. Sections were stained 5 min on drops of 2% uranyl oxalate (pH 7.0), shortly rinsed in distilled water and incubated 10 min on drops of a mixture of 1.8% (v/w) methyl cellulose/0.4% uranyl acetate (pH 4.0) on ice. Finally, grids were looped out, most of the viscous staining solution drained away and sections dried in the residual thin film which covers the grid.

The sections were analyzed using a JEM 2100Plus at 200 keV (JEOL, Japan) and a Zeiss TEM 902 at 80 keV. Labeling were controlled and imaged at same regions on three following sections on each grid.

## Suppl. Tables

**Table S 1.**
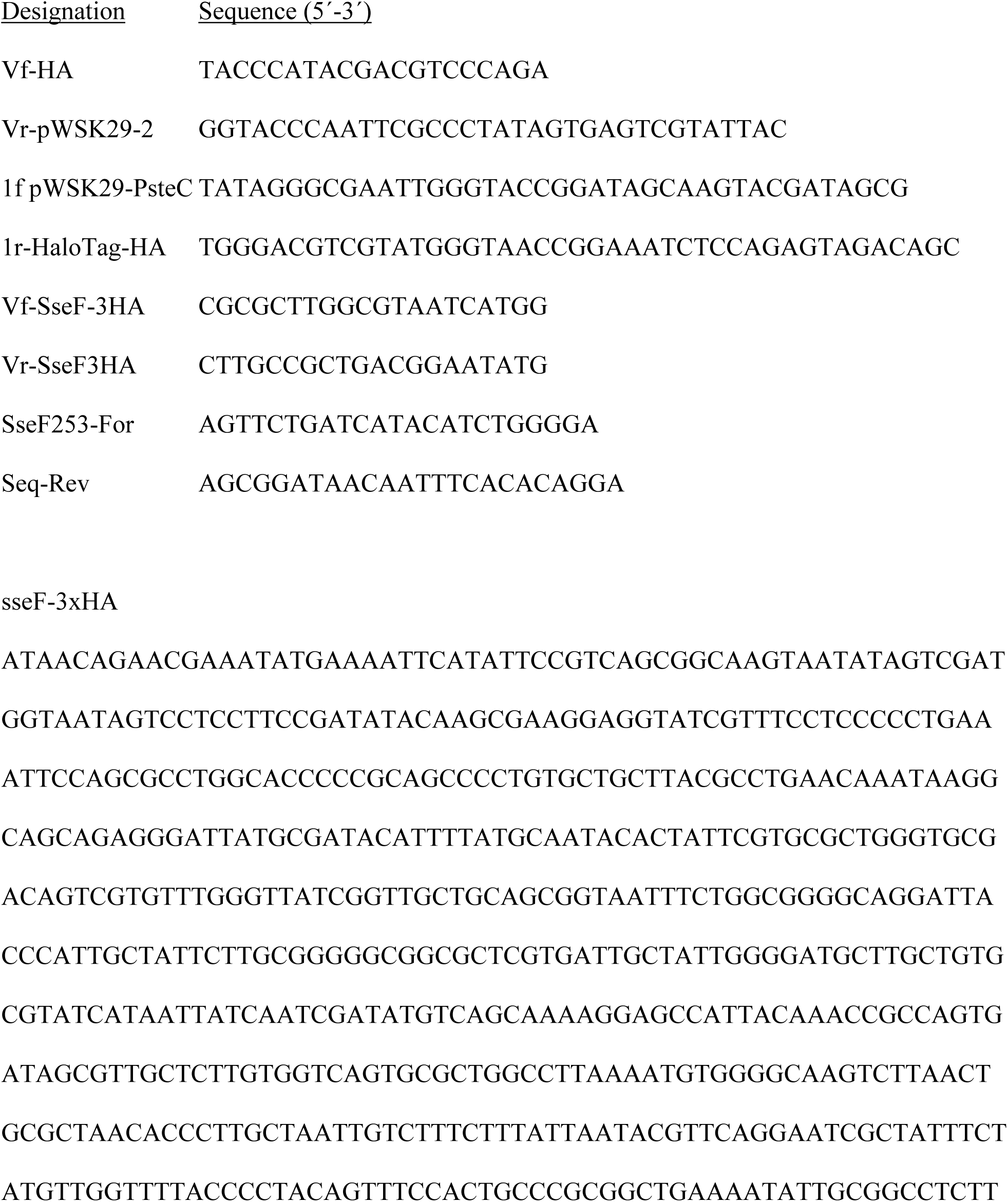
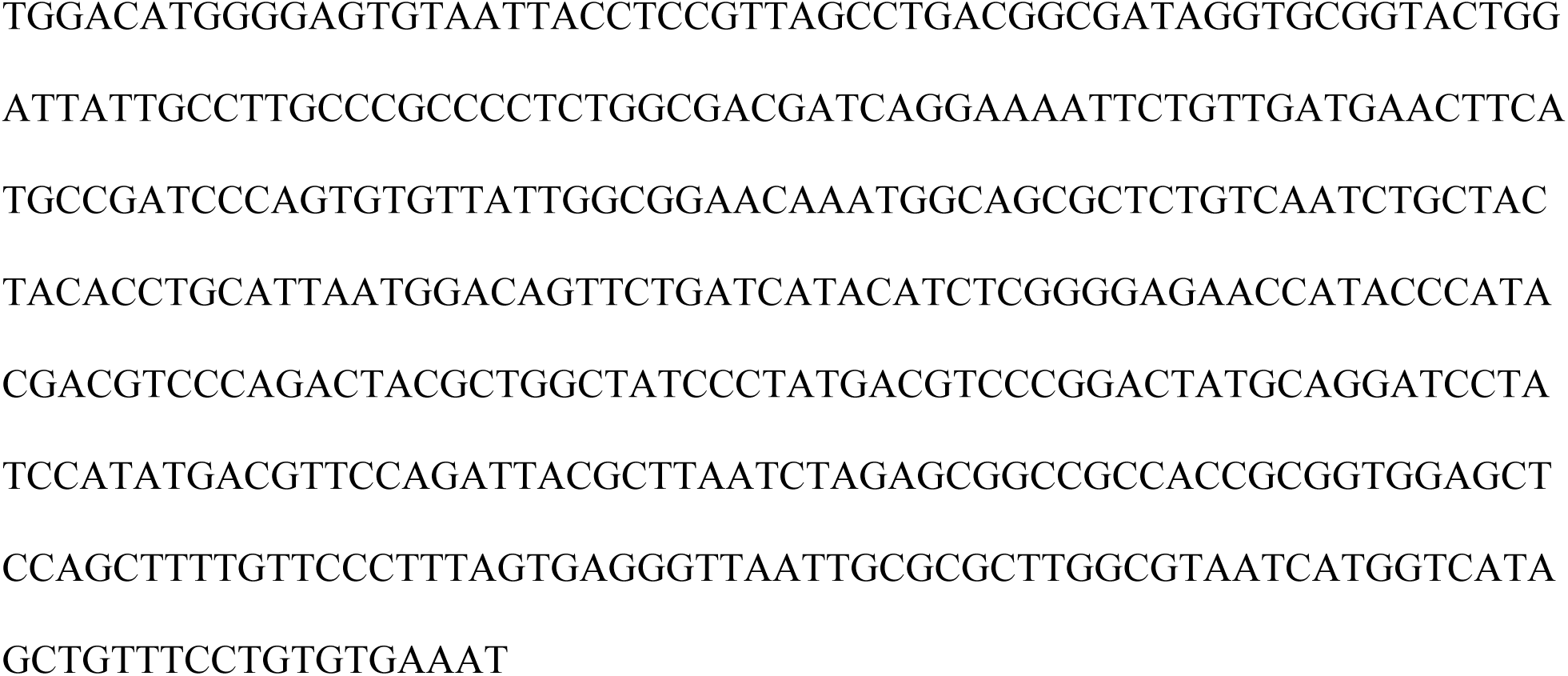
Oligonucleotides and synthetic DNA used in this study.

**Table S 2.**
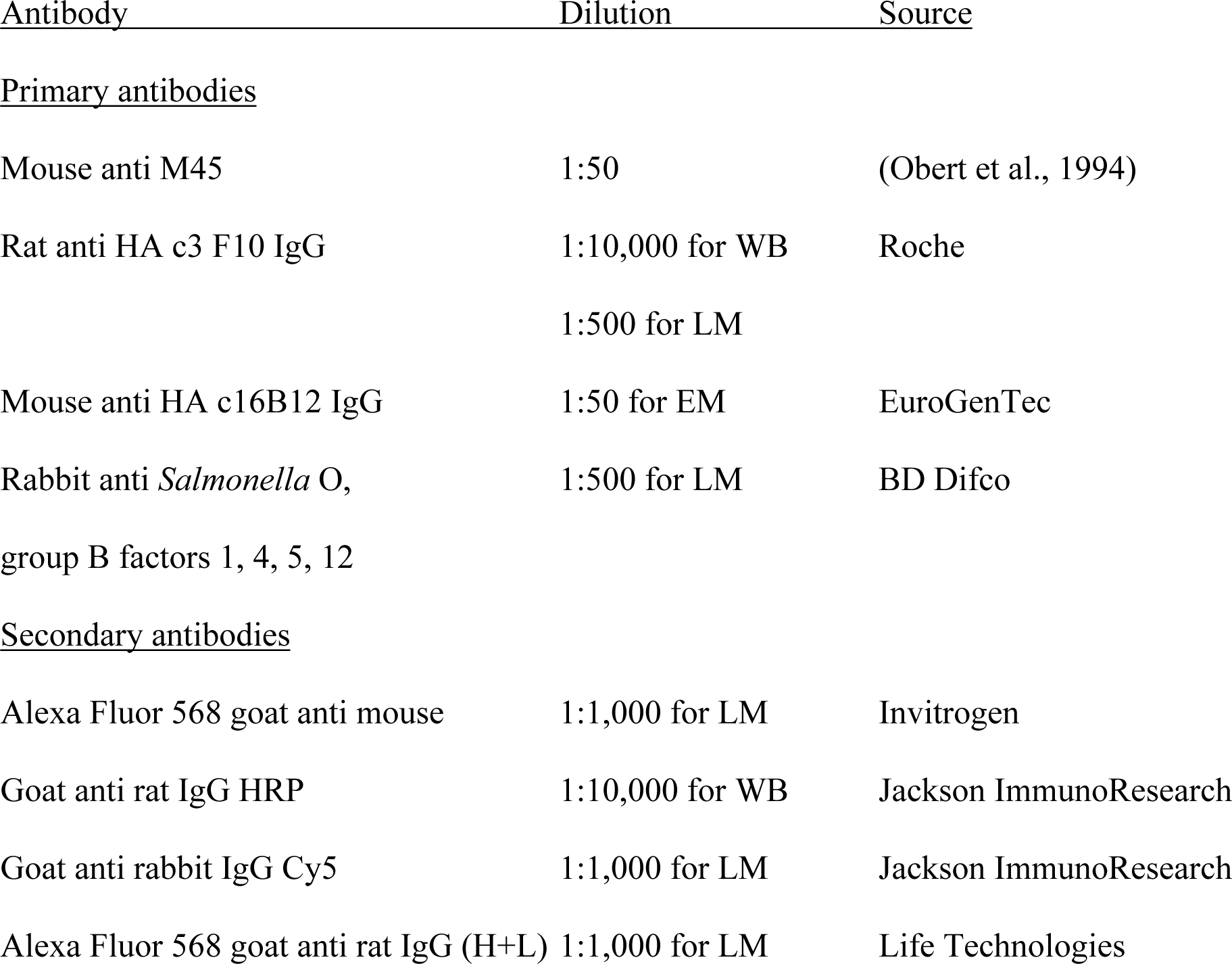
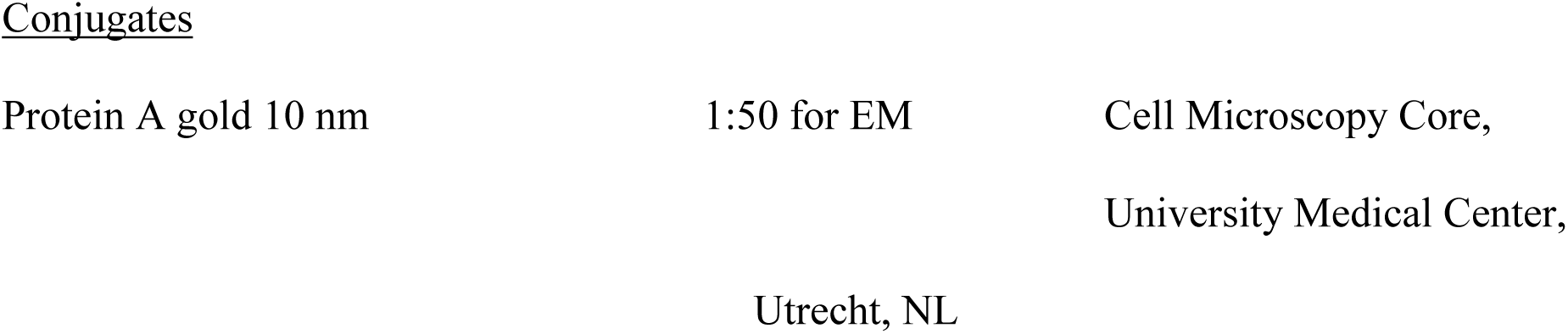
Antibodies and conjugates used in this study.

## Suppl. Figure Legends

**Fig. S 1.**
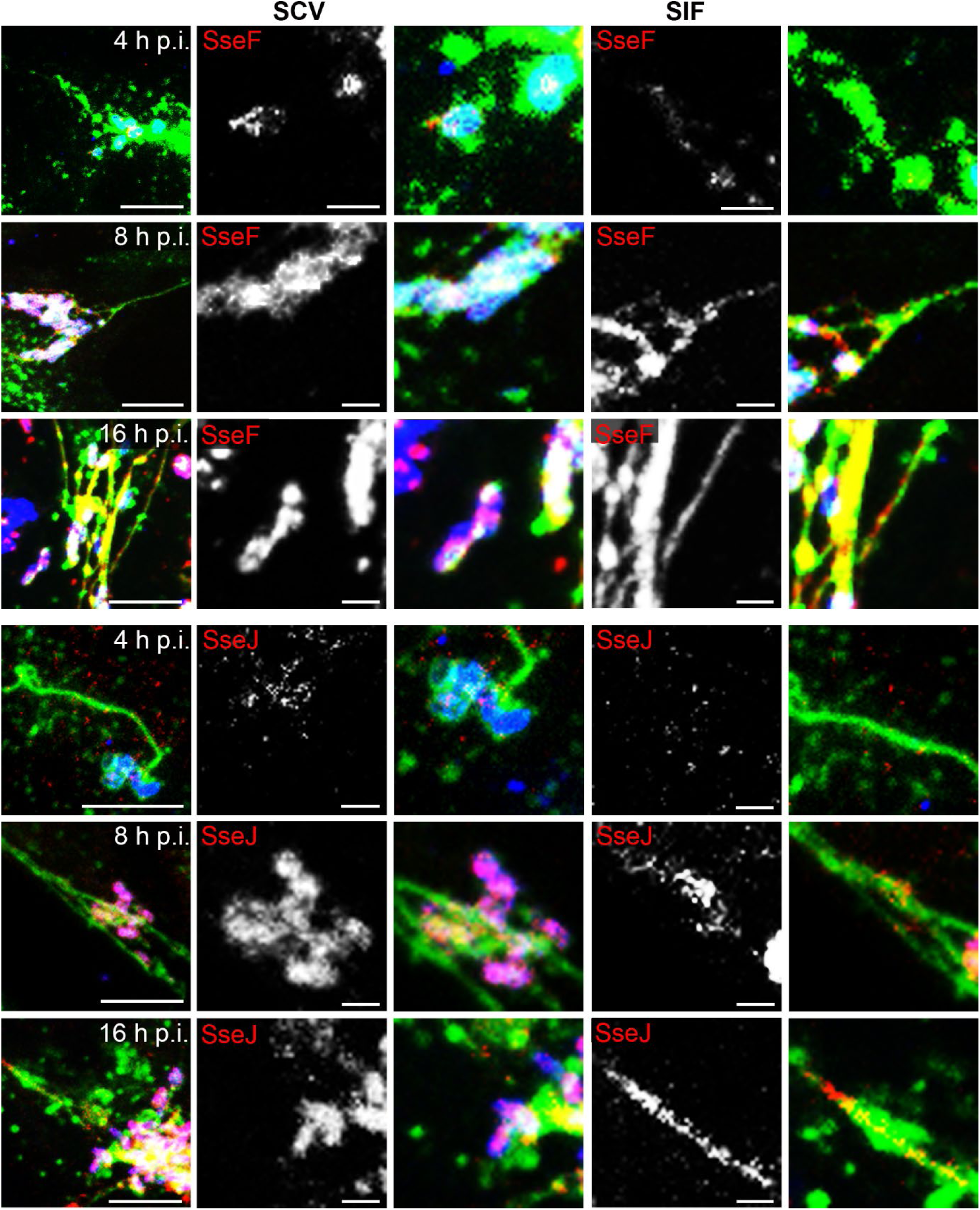
Distribution of translocated *Salmonella* SPI2-T3SS effector proteins over the course of infection. HeLa cells stably expressing LAMP1-GFP (HeLa LAMP1-GFP) were infected with STM WT expressing *sseF*::M45 or *sseJ*::M45 as indicated. At various time points after infection, cells were fixed and immunolabeled for STM (blue) and effector proteins (red). Details of SCV and SIF are shown. Scale bars: 10 and 2 µm in overview and details, respectively.

**Fig. S 2.**
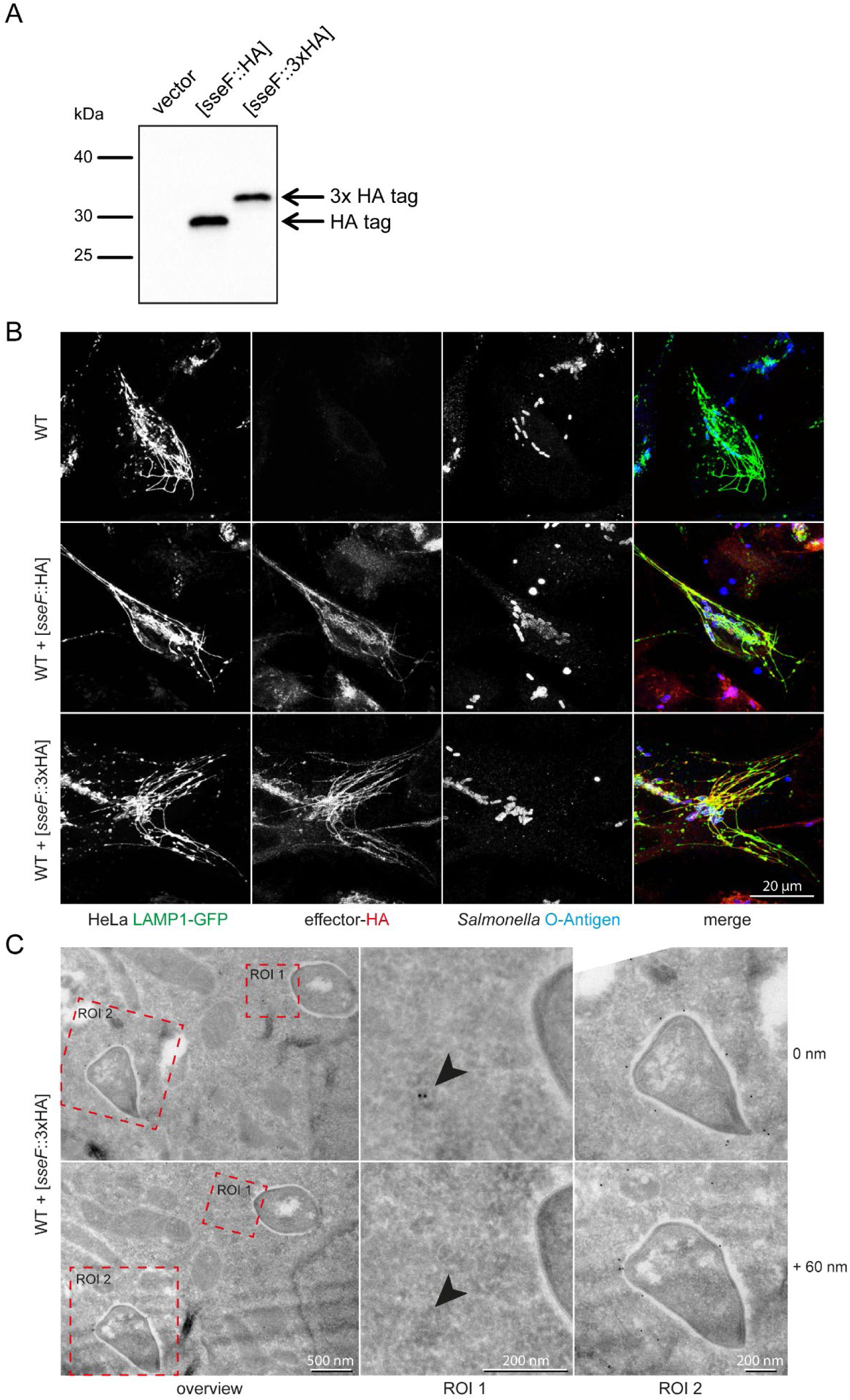
Analysis of synthesis and translocation of triple-HA-tagged effector protein SseF. **A)** Protein synthesis of triple-HA-tagged SseF after 6 h growth of subcultures in PCN (0.4) pH 5.8 medium. All plasmids were investigated in *Salmonella* wild-type background. Separation was performed by SDS-PAGE, protein was transferred onto nitrocellulose membranes and epitope-tagged proteins were detected using a primary antibody against HA tag and a secondary HRP-coupled antibody. **B**) Translocation of triple-HA-tagged SseF into stably transfected HeLa LAMP1-GFP cells. Host cells were infected, fixed 8 h p.i. and immune-stained against HA epitope tag and O-antigen. Representative cells are shown. Scale bar: 20 µm. **C**) Consecutive ultrathin sections of immunogold-labeled infected HeLa cells with STM WT expressing triple-HA-tagged SseF. Scale bars: 500 nm, 200 nm and 200 nm for overview, ROI 1 and ROI 2, respectively. Details (inserts ROI 1 and 2 in overviews) of HA-tagged SseF immunogold labeling are shown on two consecutive 60 nm thick sections. On first section (0 nm, ROI1) immunogold is located inside a vesicle (s. arrowhead), clearly indicating for a vesicular structure also proven by the following section (+60 nm, ROI 1) missing the immunogold labeling at identical region (s. arrowhead) which proves for a non-tubular structure. HA-tagged SseF immunogold labeling shown in ROI 2 on first (0 nm) and consecutive section (+60 nm) support our specific immunogold labeling for SseF.

**Fig. S 3.**
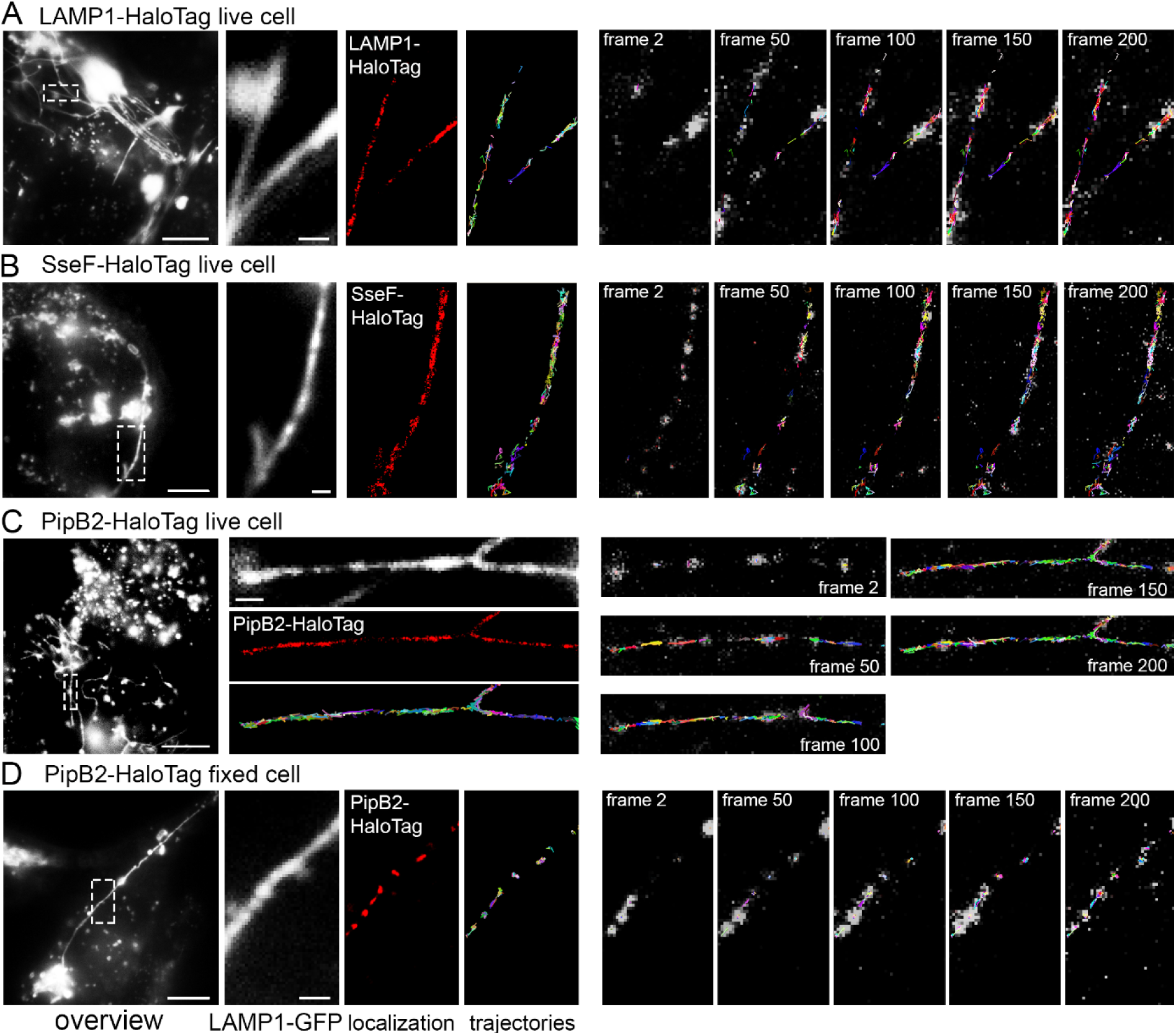
Single molecule localization (SML) and single molecule tracking (STM) of effector proteins on double-membrane SIF. HeLa LAMP1-GFP cells were infected with *Salmonella* WT or mutant strains expressing various SPI2-T3SS effector proteins fused to the HaloTag at a multiplicity of infection (MOI) of 75. For visualization of LAMP1-HaloTag, cells were transfected for expression of LAMP1::HaloTag::HA one day before infection. Following incubation for 7 h under standard cell culture conditions, live cell imaging (LCI) was performed. Labeling reactions were performed directly bevor imaging using HTL-TMR with a final concentration of 20 nM for 15 min at 37 °C. Representative SML images are shown for LAMP1-HaloTag (**A**), SseF-HaloTag (**B**), PipB2-HaloTag (**C**) in living cells, and PipB2-HaloTag in fixed cells as control (**D**). Microscopy was performed using 15% laser power at the focal plane, at 32 frames per second, SML and SMT was rendered within 200 consecutive frames. Selected frames of the TMR signal, localization and tracking are presented, also showing elapsed trajectories. Sequences of 200 frames of effector proteins or LAMP1 fused to HaloTag are shown in **Movie 3**, **Movie 4**, **Movie 5**, and **Movie 6**, corresponding to panels **A**, **B**, **C**, and **D**, respectively. Scale bars: 10 and 1 µm in overviews and details, respectively.

**Fig. S 4.**
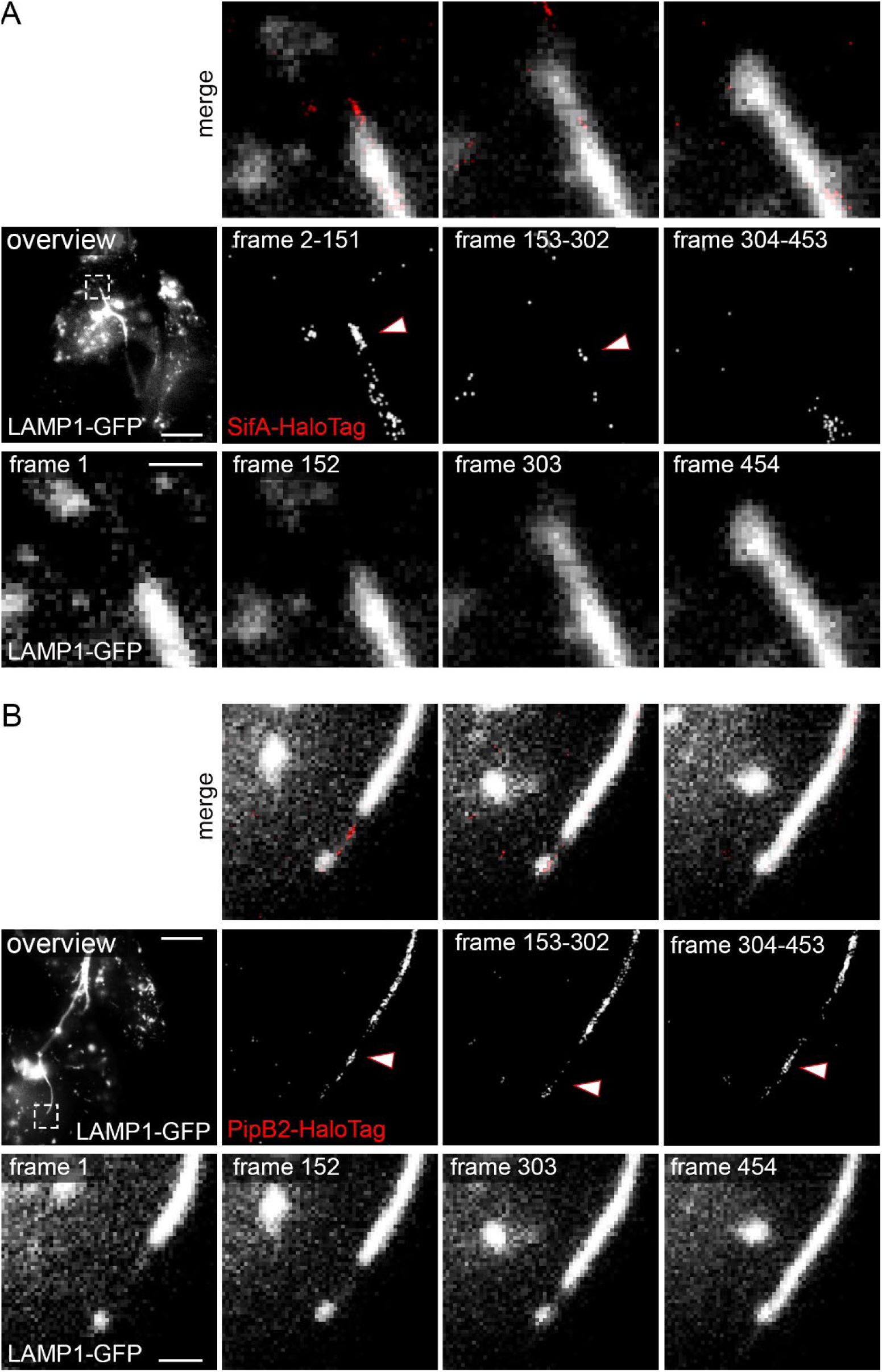
SML of effector proteins on single-membrane leading SIF. HeLa LAMP1-GFP cells were infected with STM Δ*sifA* strain expressing *sifA*::HaloTag::HA (**A**), or STM Δ*pipB2* strain expressing *pipB2*::HaloTag::HA (**B**), and labeled with HTL-TMR as described above. The transition of leading to trailing SIF was imaged with 488 nm laser excitation for one frame (frame rate: 32 frames per second) following 561 nm laser excitation for 150 frames in 4 cycles. Shown are representative SRM images acquired using 15% laser power at the focal plane, rendered from SML within each of the 150 consecutive frames. Scale bars: 10 and 1 µm in overviews and details, respectively. The sequences of 5 frames of LAMP1-GFP are shown in **Movie 10** (SifA-HaloTag) and **Movie 11** (PipB2-HaloTag).

**Fig. S 5.**
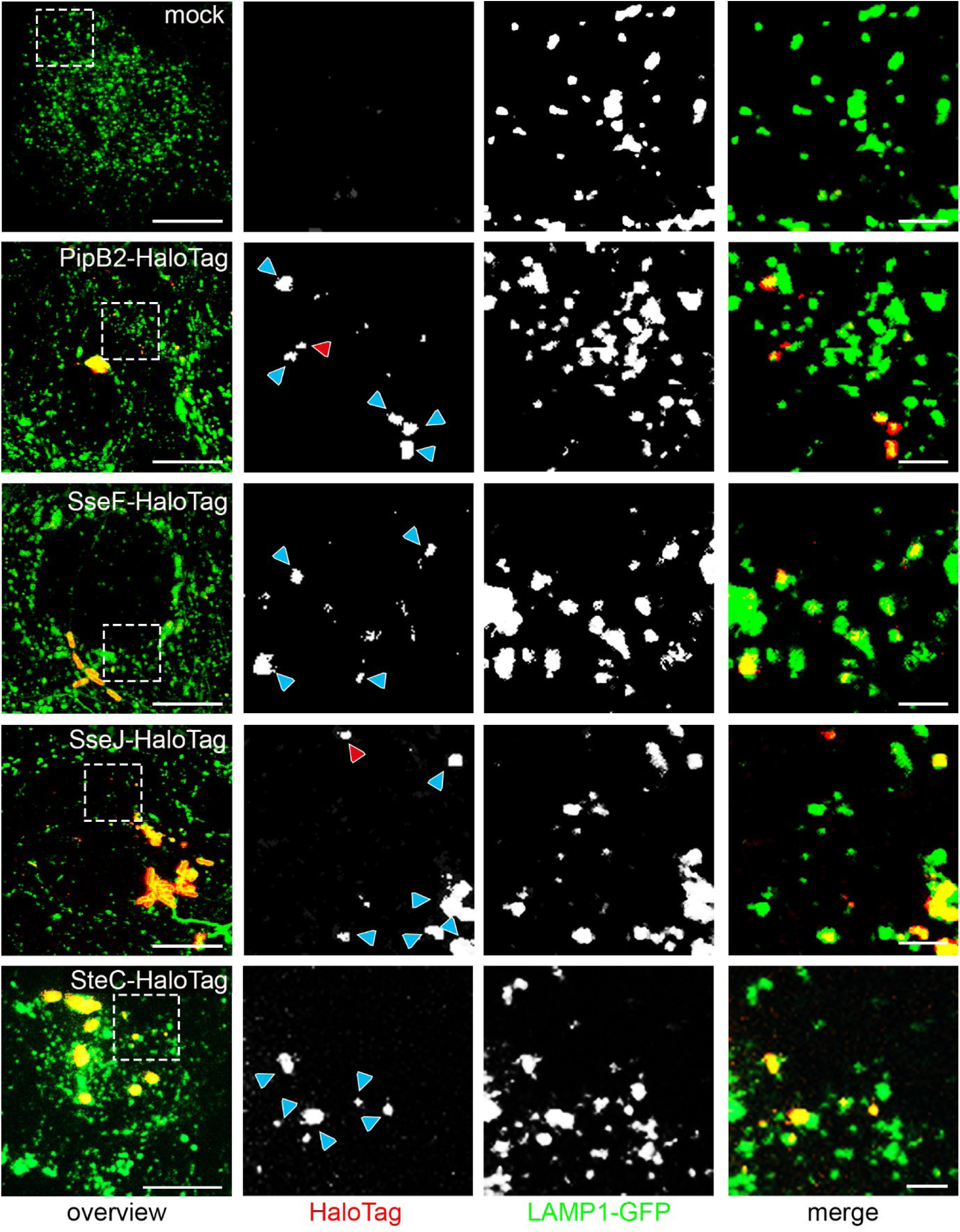
Effector protein-positive vesicles in infected HeLa LAMP1-GFP cells. HeLa LAMP1-GFP cells were infected with STM mutant strains expressing *pipB2*::HaloTag::HA, *sseF*::HaloTag::HA, *sseJ*::HaloTag::HA, or *steC*::HaloTag::HA as indicated. LCI was performed directly after cells were stained at 3.5 h p.i. with 1 µM HTL-TMR for 30 min. Shown are representative CLSM images. Blue arrowheads indicate vesicles positive for LAMP1-GFP and effector-HaloTag labeled with TMR. Red arrowheads indicate vesicles negative for LAMP1 and positive for effector-HaloTag labeled with TMR. Scale bars: 10 and 2 µm in overview and details, respectively.

**Fig. S 6.**
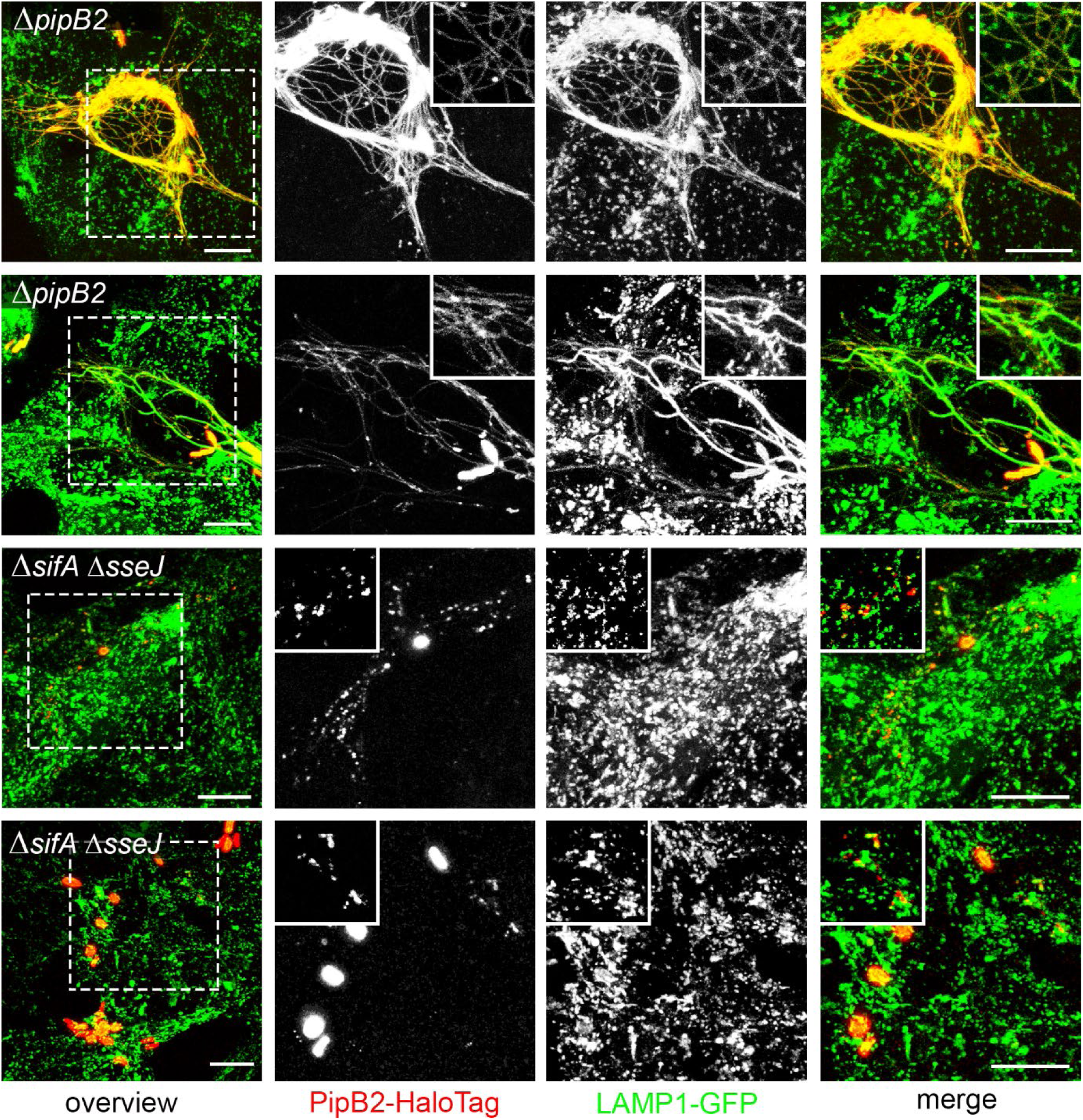
Distribution of PipB2-HaloTag in host cells infected with strains deficient in *pipB2* or Δ*sifA* Δ*sseJ*. HeLa LAMP1-GFP cells were infected with STM Δ*pipB2*, or Δ*sifA* Δ*sseJ* strains expressing *pipB2*::HaloTag::HA for 16 h. LCI was performed directly after cells were stained with HTL-TMR at a concentration of 1 µM for 30 min. Scale bars: 10 and 5 µm in overview and details, respectively.

**Fig. S 7.**
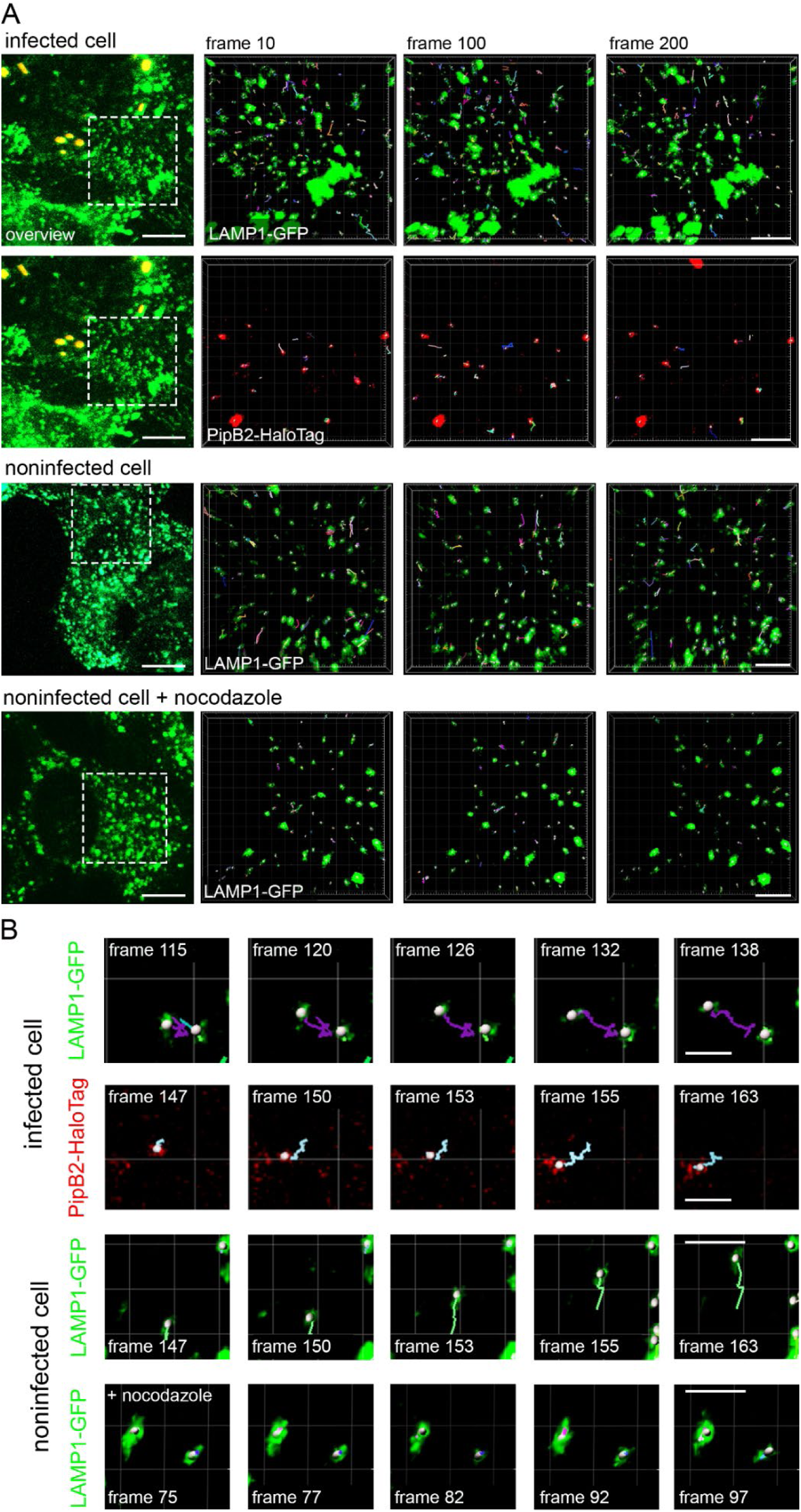
Tracking of vesicles labelled with LAMP1-GFP and PipB2-HaloTag. HeLa LAMP1-GFP cells were either not treated, infected with STM Δ*pipB2* strain expressing *pipB2*::HaloTag::HA, or treated with nocodazole to inhibit vesicle movement. **A)** Cells were imaged for 200 frames (0.39 frames/sec) using the Zeiss SD microscope and dual camera imaging in streaming mode. At least 5 cells were imaged resulting in the analysis of at least 858 trajectories/frame. Vesicle tracking analysis was done with the Imaris spot detecting tool. See corresponding **Movie 13** and **Movie 14** . **B)** Trajectories of single vesicles. Scale bars: 10 and 5 µm in overview and details, respectively.

## Movie captions

**Movie 1. Vesicular fusion of nanogold-labelled endosomes with the SCV/SIF continuum.** HeLa cells stably expressing LAMP1-GFP (green) were infected with STM WT strain constitutively expressing GFP (green). STM cells appears as high fluorescent rods, LAMP1-GFP-positive membranes are weakly fluorescent. Gold nanoparticles were prepared by conjugating 10 nm colloidal gold with BSA-rhodamine (red). Pulse/chase with Gold nanoparticles was performed 1 h p.i. for 1 h. Live cell imaging (LCI) was performed 6 h p.i. 3D projections of image stacks were generated in Imaris. Arrows indicate fusion events nanogold-labeled endosomes to a dynamic SIF/SCV continuum.

Download link: https://myshare.uni-osnabrueck.de/f/6303fcedf52a4d42b381/

**Movie 2. Single molecule tracking (SMT) of SifA-HaloTag on double-membrane SIF.** HeLa cells stably expressing LAMP1-GFP were infected with STM Δ*sifA* strain expressing *sifA*::HaloTag. LCI was performed directly after labeling with 20 nM HTL-TMR. Shown are TMR signal, localization and tracking within 200 consecutive frames (corresponding to Fig. 3A).

Download link: https://myshare.uni-osnabrueck.de/f/25e95b41fd0743a1ba69/

**Movie 3. SMT of LAMP1-HaloTag on double-membrane SIF.** HeLa LAMP1-GFP cells were transfected with LAMP1-HaloTag one day prior infection. Cells were infected with STM WT strain and LCI was done directly after labeling with 20 nM HTL-TMR. Shown are TMR signal,

localization and tracking within 200 consecutive frames (corresponding to **Fig. S 2A**).

Download link: https://myshare.uni-osnabrueck.de/f/2df9e8f60bb649fa9c82/

**Movie 4. SMT of SseF-HaloTag on double-membrane SIF.** HeLa LAMP1-GFP cells were infected with STM Δ*sseF* strain expressing *sseF*::HaloTag. LCI was done directly after labeling with 20 nM HTL-TMR. Shown are TMR signals, localization, and tracking within 200 consecutive frames (corresponding to **Fig. S 2B**).

Download link: https://myshare.uni-osnabrueck.de/f/3d369acf70ed4dc3bcc5/

**Movie 5. SMT of PipB2-HaloTag on double-membrane SIF.** HeLa LAMP1-GFP cells were infected with STM Δ*pipB2* strain expressing *pipB2*::HaloTag. LCI was done directly after labeling with 20 nM HTL-TMR. Shown are TMR signal, localization, and tracking within 200 consecutive frames (corresponding to **Fig. S 2C**).

Download link: https://myshare.uni-osnabrueck.de/f/f21d0afef4b543dca651/

**Movie 6. SMT of PipB2-HaloTag on double-membrane SIF in fixed host cells.** HeLa LAMP1-GFP cells were infected with STM Δ*pipB2* strain expressing *pipB2*::HaloTag. Cells were labeled with 20 nM HTL-TMR. Imaging was done directly after cells were fixed with 3% PFA. Shown are TMR signal, localization and tracking, within 200 consecutive frames (corresponding to **Fig. S 2D**).

Download link: https://myshare.uni-osnabrueck.de/f/570bade85d2c4868903c/

**Movie 7. SMT of LAMP1-HaloTag on single-membrane SIF.** HeLa LAMP1-GFP cells were transfected with LAMP1-HaloTag one day prior infection. Cells were infected with STM Δ*sseF* strain. LCI was done directly after labeling with 20 nM HTL-TMR. Shown are TMR signal, localization, and tracking within 200 consecutive frames (corresponding to Fig. 4A).

Download link: https://myshare.uni-osnabrueck.de/f/1c42b10e1f414ab0a3b3/

**Movie 8. SMT of SifA-HaloTag on single-membrane SIF.** HeLa LAMP1-GFP cells were infected with STM Δ*sseF* Δ*sifA* strain expressing *sifA*::HaloTag. LCI was performed directly after labeling with 20 nM HTL-TMR. Shown are TMR signal, localization, and tracking within 200 consecutive frames (corresponding to Fig. 4B).

Download link: https://myshare.uni-osnabrueck.de/f/919d7993597e40948b9b/

**Movie 9. Localization of SseF-HaloTag during transition of leading to trailing SIF.** HeLa LAMP1-GFP cells were infected with STM Δ*sseF* strain expressing *sseF*::HaloTag and labeled with HTL-TMR. The transition of leading to trailing SIF was imaged with 488 nm laser excitation for one frame, following 561 nm laser excitation for 150 frames in 4 cycles. Shown are 5 frames of 488 nm excitation (corresponding to Fig. 4E).

Download link: https://myshare.uni-osnabrueck.de/f/2cb53caa8c81451c89a4/

**Movie 10. Localization of SifA-HaloTag during transition of leading to trailing SIF.** HeLa LAMP1-GFP cells were infected with STM Δ*sifA* strain expressing *sifA*::HaloTag and labeled with HTL-TMR. The transition of leading to trailing SIF was imaged with 488 nm laser excitation for one frame, following 561 nm laser excitation for 150 frames in 4 cycles. Shown are 5 frames of 488 nm excitation (corresponding to **Fig. S 3A**).

Download link: https://myshare.uni-osnabrueck.de/f/4851e1aaa09b437aa0a4/

**Movie 11. Localization of PipB2-HaloTag during transition of leading to trailing SIF.** HeLa LAMP1-GFP cells were infected with STM Δ*pipB2* strain expressing *pipB2*::HaloTag and labeled with HTL-TMR. The transition of leading to trailing SIF was imaged with 488 nm laser excitation for one frame, following 561 nm laser excitation for 150 frames in 4 cycles. Shown are 5 frames of 488 nm excitation (corresponding to **Fig. S 3A**).

Download link: https://myshare.uni-osnabrueck.de/f/777f93142ae7459ca5fc/

**Movie 12. Co-tracking of LAMP1-GFP- and PipB2-HaloTag-positive vesicles in infected host cells.** HeLa LAMP1-GFP cells were infected with STM Δ*pipB2* strain expressing *pipB2*::HaloTag. LAMP1-positive- and PipB2-HaloTag-positive vesicles were imaged and vesicle tracking analysis was done with Imaris spot detection tool. Shown are 200 frames of co-tracking of vesicles positive for LAMP1 and PipB2 co-tracking (corresponding to Fig. 6A).

Download link: https://myshare.uni-osnabrueck.de/f/095efdca37b14916ad6a/

**Movie 13. Tracking of LAMP1-GFP- and PipB2-HaloTag-positive vesicles in STM-infected host cells.** HeLa LAMP1-GFP cells were infected with STM Δ*pipB2* strain expressing *pipB2*::HaloTag. LAMP1-positive and PipB2-positive vesicles were tracked with the Imaris spot detection tool in individual cells over 200 frames (corresponding to Fig. 6A).

Download link: https://myshare.uni-osnabrueck.de/f/2bc6b6801d8240449e47/

**Movie 14. Tracking of LAMP1-GFP-positive vesicles in non-infected and nocodazole-treated cells.** HeLa LAMP1-GFP cells either non-treated or treated with nocodazole. LAMP1-positive vesicles were tracked with the Imaris spot detection tool in individual cells over 200 frames (corresponding to Fig. 6A).

Download link: https://myshare.uni-osnabrueck.de/f/a633c24d92f24442a4a2/

**Movie 15. PipB2-HaloTag distribution in STM-infected cells.** HeLa LAMP1-GFP cells were infected with STM Δ*pipB2* strain expressing *pipB2*::HaloTag. Time-lapse imaging was performed directly after cells were stained with HTL-TMR (1 µM). Cells were imaged every 30 min starting 5 h p.i. until 12 h p.i. (corresponding to Fig. 7A).

Download link: https://myshare.uni-osnabrueck.de/f/5e80ad49973a43e8a9d5/

**Movie 16. Effect of nocodazole on PipB2-HaloTag distribution in STM-infected cells** HeLa LAMP1-GFP cells were infected with STM Δ*pipB2* strain expressing *pipB2*::HaloTag. Cells were treated with nocodazole (2 h p.i.). Time-lapse imaging was performed directly after cells were stained with HTL-TMR (1 µM) and inhibitor was removed. Cells were imaged every 30 min starting 5 h p.i. until 12 h p.i. (corresponding to Fig. 7AB).

Download link: https://myshare.uni-osnabrueck.de/f/55645ebb2c4a4b89a1aa/

## Notes

### Competing Interest Statement

The authors have declared no competing interest.

